# Tau forms oligomeric complexes on microtubules that are distinct from pathological oligomers in disease

**DOI:** 10.1101/2020.07.07.192146

**Authors:** M. T. Gyparaki, A. Arab, E. M. Sorokina, A. N. Santiago-Ruiz, C. H. Bohrer, J. Xiao, M. Lakadamyali

## Abstract

Tau is a microtubule-associated protein, which promotes neuronal microtubule assembly and stability. Accumulation of tau into insoluble aggregates known as neurofibrillary tangles (NFTs) is a pathological hallmark of several neurodegenerative diseases. The current hypothesis is that small, soluble oligomeric tau species preceding NFT formation cause toxicity. However, thus far visualizing the spatial distribution of tau monomers and oligomers inside cells under physiological or pathological conditions has not been possible. Here, using single molecule localization microscopy (SMLM), we show that, *in vivo*, tau forms small oligomers on microtubules under physiological conditions. These physiological oligomers are distinct from those found in cells exhibiting tau aggregation and could be pre-cursors of aggregated tau in pathology. Further, using an unsupervised shape classification algorithm that we developed, we show that different tau phosphorylation states are associated with distinct tau aggregate species. Our work elucidates tau’s nanoscale composition under physiological and pathological conditions *in vivo*.

## Introduction

Tau is a microtubule-associated protein (MAP), mainly expressed in neurons in humans. Tau is an intrinsically disordered protein, which facilitates the assembly and stability of neuronal microtubules. Immunofluorescence labeling showed that tau is highly abundant in neuronal axons and shows a graded distribution with higher concentration towards the axon terminal^1^. The first near-atomic model of tau bound to microtubules obtained using cryoelectron microscopy (cryo-EM) and computational modeling revealed that tau binds along the microtubule protofilament stabilizing the interface between tubulin dimers^2^. Tau has also been shown to undergo liquid-liquid phase separation *in vitro* and tau condensates have been proposed to facilitate microtubule polymerization due to their ability to sequester soluble tubulin^3^. *In vitro* reconstitution experiments further showed that tau forms liquid condensates that are several microns in size on taxol-stabilized microtubules and these tau “islands” regulate microtubule severing and microtubule-motor protein interactions^4,5^. However, despite progress, the distribution of tau on microtubules in intact cells and the physiological relevance of tau condensates are unclear.

Under pathological conditions, tau becomes hyper-phosphorylated and detaches from microtubules leading to misfolding and formation of tau aggregates in the cytosol^6^. Accumulation of tau into insoluble aggregates known as neurofibrillary tangles (NFTs) is a hallmark of several neurodegenerative diseases such as Alzheimer’s Disease (AD) and Frontotemporal Dementia with Parkinsonism linked to chromosome 17 (FTDP-17), which are collectively known as tauopathies. Liquid phase separation has also been suggested to play a role in pathological tau aggregation as aggregation prone tau mutants have an increased propensity to undergo liquid phase separation^3^. While NFTs have traditionally been considered the pathological tau species, recent evidence from animal models has shown that neurodegeneration in terms of synaptic dysfunction and behavioral abnormalities can exist without the presence of NFTs^7,8^. The current hypothesis is that earlier stages of tau aggregation initiate pathology. In particular, small, soluble tau oligomeric species preceding the formation of NFTs could be the cause of loss of tau function and toxicity. Soluble tau oligomers have been isolated from homogenized AD brains and detected on SDS-PAGE gels^9^. These tau oligomers have also been implicated in synaptic loss in transgenic mice expressing wild-type human tau^10,11,12^. Moreover, injection of tau oligomers rather than monomers or fibrils into the brain of wild-type mice was sufficient to produce cognitive and synaptic abnormalities^13,14^. Tau oligomers can lengthen and adopt a β-sheet conformation, which renders them detergent-insoluble. Such oligomers have granular appearance under atomic force microscopy (AFM)^15^. Despite their *in vitro* characterization, to date, there have not been studies visualizing and characterizing the formation of these small oligomers in intact cells due to the lack of quantitative, high-resolution tools. The lack of sensitive tools that enable visualizing the spatial distribution and quantifying the stoichiometry of tau complexes within native and diseased neurons prevents progress in studying the mechanisms that lead to pathological tau oligomerization and the impact of early stages of tau aggregation on cellular processes.

Here, we have used quantitative super-resolution microscopy to visualize and determine the nanoscale organization of tau under physiological conditions in engineered cell models^16^ as well as in neurons. In addition, using an engineered cell model (Clones 4.0 and 4.1) expressing tau harboring the FTDP-17 mutation P301L to mimic tau aggregation in disease^16^, we characterized the nature of tau aggregates including oligomeric tau species. Surprisingly, our results show that, in engineered cell lines, tau forms a patchy distribution consisting of nano-clusters along the microtubule under physiological conditions. Tau nano-clusters mainly correspond to monomeric, dimeric and trimeric tau complexes. The nanoscale distribution of microtubule-associated tau in these engineered cell lines is not strongly dependent on tau expression levels and resembles the nanoscale distribution of microtubule-associated tau in hippocampal neurons isolated from rats. In pathological conditions (aggregated tau Clone 4.1), tau forms a diverse range of aggregate species including small oligomeric tau assemblies, small fibrillary structures, branched fibrils and large plaque-like structures resembling NFTs. The pathological oligomeric species are partially microtubule-associated and are distinct from the physiological ones as they are larger, containing more tau molecules. Using antibodies that recognize different phosphorylation states of tau and an unsupervised machine learning based shape classification method, we further show that different phosphorylation states associate with distinct higher order tau aggregate species in pathology. To our knowledge, this is the first characterization of tau oligomers on microtubules in intact cells and the first detailed characterization of tau aggregates with nanoscale resolution in a cell model of FTDP-17.

## Results

### Microtubule-associated tau is partially oligomeric under physiological conditions

To determine how tau is distributed under physiological conditions, we performed single molecule localization microscopy (SMLM) in two different engineered cell models: one that constitutively expresses wild-type (WT) tau fused to green fluorescent protein, GFP, (GFP-tau) and a second inducible cell line that expresses GFP-tagged tau harboring a FTDP-17 mutation (P301L) (GFP-P301L-tau) under the control of doxycycline (Dox) (Clone 4.0). The pattern of GFP expression in both engineered cell lines resembles the microtubule network (Figure S1A), indicating that tau is predominantly microtubule associated in both cases as expected. Hence, GFP fusion does not interfere with tau’s microtubule binding ability. Since these engineered cell lines do not normally express tau, we asked whether when tau expression is induced by Dox in Clone 4.0, the binding of tau to the microtubules impacts the microtubule network. To address this question, we imaged the microtubule network in Clone 4.0 before and after Dox induction using SMLM. As expected, tau expression led to increased microtubule density and microtubule bundling, in line with the known functions of tau in facilitating microtubule polymerization and bundling (Fig S1B, top and middle; Fig S1C, first two plots)^17,18^. These results further support that GFP fusion did not disrupt tau’s function.

We next labeled GFP-tau and GFP-P301L-tau Clone 4.0 with a GFP nanobody conjugated with a photoswitchable fluorescent dye (AF647) (Fig.1A) to image tau’s nanoscale distribution with SMLM. To ensure that fixation did not perturb tau localization, we tested two commonly used fixation protocols using either aldehyde based fixation (paraformaldehyde, PFA) or alcohol based fixation (methanol) (see Methods). PFA fixation led to mis-localization of tau and loss of “microtubule-like” staining visible in live cells (Figure S2A) in line with previous reports^19,20^. Methanol fixation, on the other hand preserved the microtubule localization of tau (Figure S2A). Indeed, when we determined the fluorescence intensity of microtubule associated GFP-tau in the same cells before and after methanol fixation, the fluorescence intensity was only reduced by a small amount (Mean Intensity: 871.2 AU before and 750.1 AU after fixation), suggesting that the majority of tau remains microtubule bound after fixation (Figure S2B). Hence, we used methanol fixation for visualizing tau’s nano-scale organization. SMLM images in both cell lines revealed that both WT tau and P301L-tau are not uniformly distributed on the microtubule, unlike α-tubulin (Fig 1A and Fig S2C) but instead form nano-clusters. We segmented these nano-clusters using Voronoi tessellation (^21,22^ and Methods, Fig 1A and Fig S2D) and quantified the number of localizations per nano-cluster (corresponding to the number of times an AF647 fluorophore was localized) as well as the nano-cluster area (Fig 1C, Fig S2E), obtaining a broad distribution in both cases. These parameters were similar for GFP-tau and GFP-P301L-tau Clone 4.0 despite the large differences in tau expression level in the two cell lines (6-fold) (Fig S2F), suggesting that tau distribution on the microtubule does not strongly depend on expression levels. Since tau distribution was similar between GFP-tau and GFP-P301L-tau, we used the GFP-P301L-tau cell line for subsequent analysis.

**Fig. 1.**
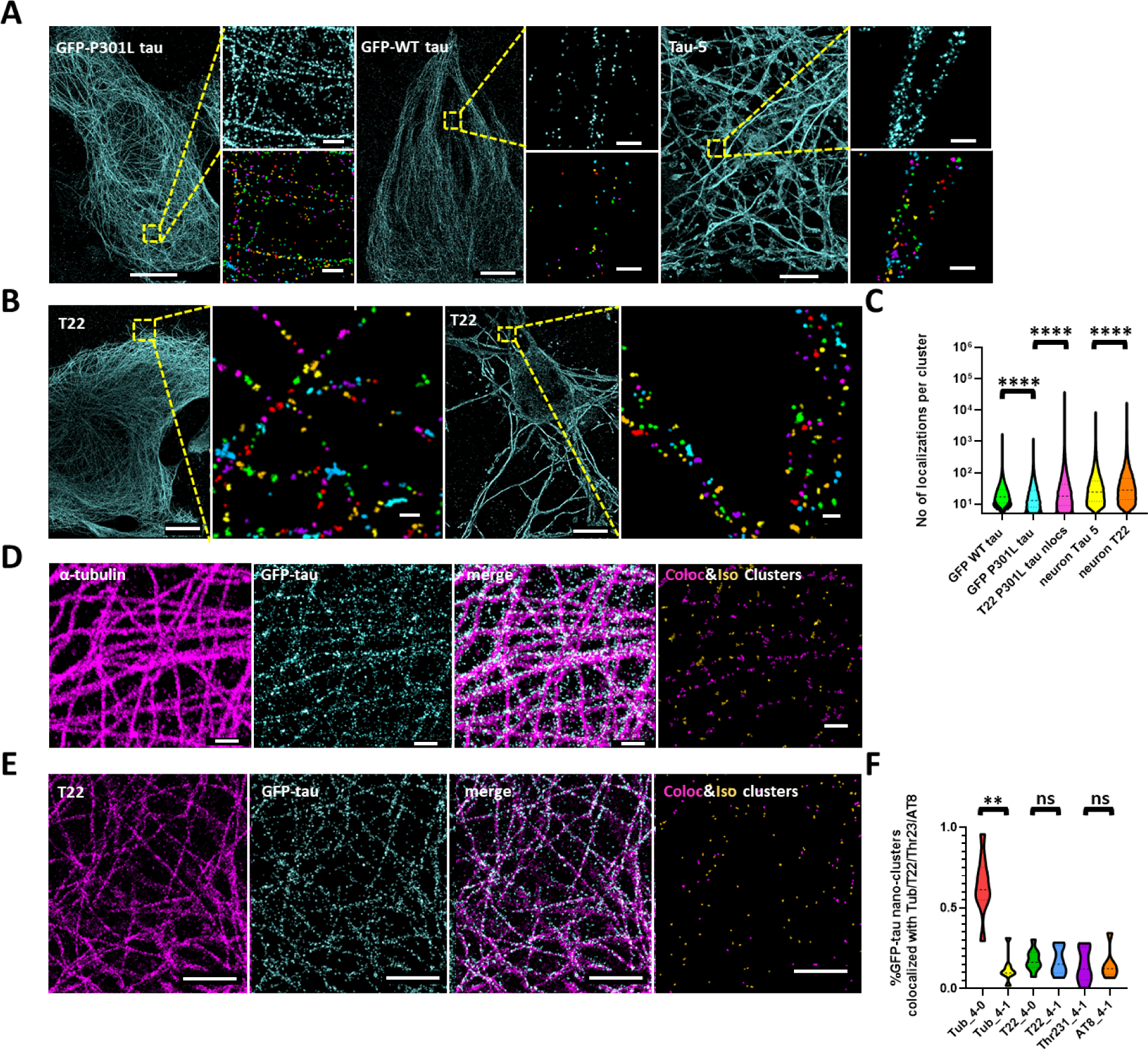
Microtubule-associated tau is partially oligomeric under physiological conditions. **A**. Super-resolution images of tau in GFP-P301L tau (Clone 4.0) and GFP-WT tau cells stained with GFP nanobody conjugated to AlexaFluor 647 as well as in neurons stained with Tau-5 antibody (from left to right). Zoomed in regions of all images are shown together with their corresponding Voronoi segmentation. Segmented images are pseudo color-coded with different colors corresponding to different segmented nano-clusters. Scale bars in zoom out images are 10um and 200nm for zoom in images. **B**. Super-resolution images of oligomeric tau detected by T22 antibody in GFP-P301L tau cells (Clone 4.0) and in neurons (from left to right). Zoomed in regions show pseudo color-coded Voronoi segmentation of nano-clusters. Scale bars in zoom out images are 10um and 200nm for zoom in images. **C**. Violin plots showing the number of localizations per nano-cluster segmented with Voronoi segmentation in the different cell lines used in this study. The dashed lines indicate the median and the dotted lines indicate the 25^th^ and 75^th^ percentile (GFP-WT tau: n=15 cells, n=2 experiments, GFP-P301L tau: n=15 cells, n= 3 experiments, T22 P301L tau: n=19 cells, n=3 experiments, neuron Tau 5: n=3 cells, neuron T22: n=3 cells). ****, *p*<0.0001. **D**. Two-color super-resolution images of α-tubulin (magenta), total tau (cyan) and overlay in GFP-P301L tau cells (Clone 4.0). The results of the co-localization analysis are shown in which tau nano-clusters co-localized with α-tubulin are color-coded in magenta and isolated tau nano-clusters are shown in yellow. Scale bars are 500nm. **E**. Two-color super-resolution images of oligomeric tau detected by T22 (magenta), total tau (cyan) and overlay in GFP-P301L tau cells (Clone 4.0). The results of the co-localization analysis are shown in which tau nano-clusters co-localized with T22 are color-coded in magenta and isolated tau nano-clusters are shown in yellow. Scale bares are 2um. **F**. Violin plots showing the % of tau nano-clusters co-localized with α-tubulin, T22, Thr231 and AT8 in GFP-P301L-tau cells (Clone 4.0 and Clone 4.1) (Tub 4-0: n=12 cells, n=3 experiments, Tub 4-1: n=9 cells, n=3 experiments, T22 4-0: n=9 cells, n=3 experiments, T22 4-1: n=6 cells, n=2 experiments, Thr231 4-1: n=6 cells, n=2 experiments, AT8: n=6 cells, n=3 experiments). **, *p*<0.01. ns, non-significant.

To determine the monomeric versus oligomeric nature of tau within these nano-clusters, we used a number of different approaches. In principle, the number of localizations per nano-cluster in SMLM images is proportional to the number of proteins. However, repeated fluorophore blinking and the fact that the labeling ratio between the protein and the fluorophore is often not one-to-one typically leads to over-counting artifacts^23^. First, to exclude the possibility that these are artificial nano-clusters resulting from repeated blinking of the same fluorophore, we used Distance Distribution Correction (DDC)^24^ to eliminate multiple localizations from the same molecule. Although DDC narrowed down the distribution of GFP-tau nano-cluster areas and the number of localizations per cluster (Fig S3A-C), the distributions still remained broad. Importantly, DDC correction did not eliminate the presence of nano-clusters, suggesting that these are not artificial nano-clusters due to repeated fluorophore blinking. Second, we used a commercial oligomeric tau antibody (T22) to specifically stain oligomeric tau. SMLM images of the T22 labeled cells revealed discontinuous microtubule-like pattern similar to GFP-tau and GFP-P301L tau suggesting that the oligomeric T22 antibody binds to microtubule-associated tau (Fig.1B).

Taken together, our analyses suggested that observed tau nano-clusters contain oligomeric tau molecules under physiological conditions. To estimate the number of tau molecules in these nano-clusters, we next used a method that we previously developed to estimate the stoichiometry of proteins in super-resolution images using the number of localizations distribution from the monomeric protein images as a calibration^25^. To determine the number of localizations distribution corresponding to the monomeric protein, we performed a 100-fold dilution of the GFP nanobody to sparsely label individual tau molecules instead of saturating all GFP-epitopes. This approach has previously been successfully used to calibrate antibody labeled receptors to determine their stoichiometry within synapses^26^. The resulting images under dilute labeling conditions showed “nano-clusters” that were much sparser than those in densely labeled cells (Fig S3D). “Nano-clusters” under dilute labeling conditions also contained significantly lower number of localizations compared to those in densely labeled cells (Fig S3E). We assumed that the “nano-clusters” under sparse labeling conditions thus mainly corresponded to monomeric tau and appeared as “nano-clusters” due to repeated blinking of the same fluorophore. Indeed, DDC correction eliminated the appearance of “nano-clusters” (Fig S3F) and Voronoi segmentation of DDC corrected images did not detect any “nano-clusters” under dilute labeling conditions. Using the distribution of the number of localizations per “nano-cluster” prior to DDC correction under dilute labeling conditions as a calibration (see Methods), we determined that tau nano-clusters in the densely labeled GFP-P301L-tau cells contain approximately ∼60% monomers, ∼20% dimers and ∼20% trimers (Fig S3G,H). Tau nano-clusters in the GFP-WT-tau line consisted of ∼60% monomers, ∼25% dimers and ∼15% trimers, which is very similar to the percentages determined in the GFP-P301L-tau cell line (Fig S3I).

Finally, to confirm that the nano-clusters are not due to overexpression of GFP-fusion protein in the engineered cell lines, we stained endogenous tau in hippocampal neurons with a total tau antibody, Tau-5 (Fig 1A) as well as T22 (Fig 1B) and obtained similar results in which tau formed nano-clusters containing a similarly wide distribution of number of localizations as in cells (Fig 1C). We repeated the dilute labeling experiments by lowering the concentration of the primary Tau-5 antibody by 100-fold while keeping the secondary antibody concentration at the same level in neurons. Under dilute primary antibody labeling conditions, we obtained sparse nano-clusters containing significantly fewer localizations and likely corresponding to monomeric tau similar to the nanobody experiments (Figure S3J). These results indicate that tau oligomers are also present in physiological conditions in cultured neurons and are not an artefact of GFP-fusion.

To further determine the percentage of oligomeric tau present in cells as well as the amount of soluble versus microtubule-associated tau, we labeled tau together with α-tubulin or with T22 and carried out two-color SMLM (Fig. 1D-E). We used the images of microtubules or of oligomeric tau labeled with T22 antibody as a mask and categorized the tau nano-clusters into two populations: those that co-localize with the reference mask and those that are isolated (see Methods) (Fig. 1D,E). Approximately 61±19% of the GFP-tau nano-clusters were microtubule-associated (Fig. 1F). Similarly, ∼20±7% of GFP-tau nano-clusters co-localized with T22 stained tau oligomers (Fig. 1F). A positive control experiment in which we labeled GFP-tau with nanobodies in two colors and carried out co-localization analysis showed approximately 60±13% co-localization under conditions in which we expect full co-localization (Fig. S4A), consistent with previous 2-color super-resolution results^27^. Hence, our two color imaging and co-localization analysis underestimates the true extent of co-localization by ∼1.7-fold, suggesting that tau is almost exclusively microtubule associated in this cell line and ∼34% of tau co-localizes with T22. The latter number is in line with our calibration and stoichiometry estimation showing ∼40% tau dimers/trimers.

Based on these results, we concluded that under physiological conditions tau is primarily microtubule associated and a proportion of tau that is associated with microtubules forms dimeric and trimeric complexes.

### Tau oligomers in cells exhibiting tau aggregation are distinct from physiological tau oligomers

To determine the nanoscale distribution and composition of tau aggregates under pathological conditions, we used an engineered cell line model of FTDP-17: Clone 4.1 GFP-P301L-tau, which is a sub-clone of the Clone 4.0 GFP-P301L-tau cell line. Sub-clone 4.1 was obtained after transfecting the parent (4.0) cell line with tau fibrils to induce tau aggregation and selected because it stably maintains tau aggregates when cultured in the presence of Dox^16^. Previous work using this sub-clone showed that it recapitulates tau aggregation seen in disease states, such as formation of large tau inclusions resembling NFTs^16^. Further, using this cell line it was shown that lysosomal degradation plays a key role in clearing tau aggregates^16^. These previous results suggest that Clone 4.1 is a good model system for pathological tau aggregation and for gaining new insights into mechanisms of tau aggregation and clearance *in vivo*. SMLM images of tau in Clone 4.1 were dramatically different from those of Clone 4.0 (Fig 2A). While a small population of microtubule-associated tau remained, there was a large population of cytosolic tau aggregates having diverse sizes and shapes. The resulting Voronoi segmented objects ranged from small nano-clusters, similar to those observed in the physiological conditions, to fibrillary structures and large conglomerate aggregates that are reminiscent of NFTs in terms of their size (Fig 2A). The number of localizations per segmented object and the area of the segmented objects were very broad and on average higher compared to the nano-clusters in Clone 4.0, in line with the visual impression of the presence of a wide range of aggregate structures (Fig 2B, Fig S4B).

**Fig. 2.**
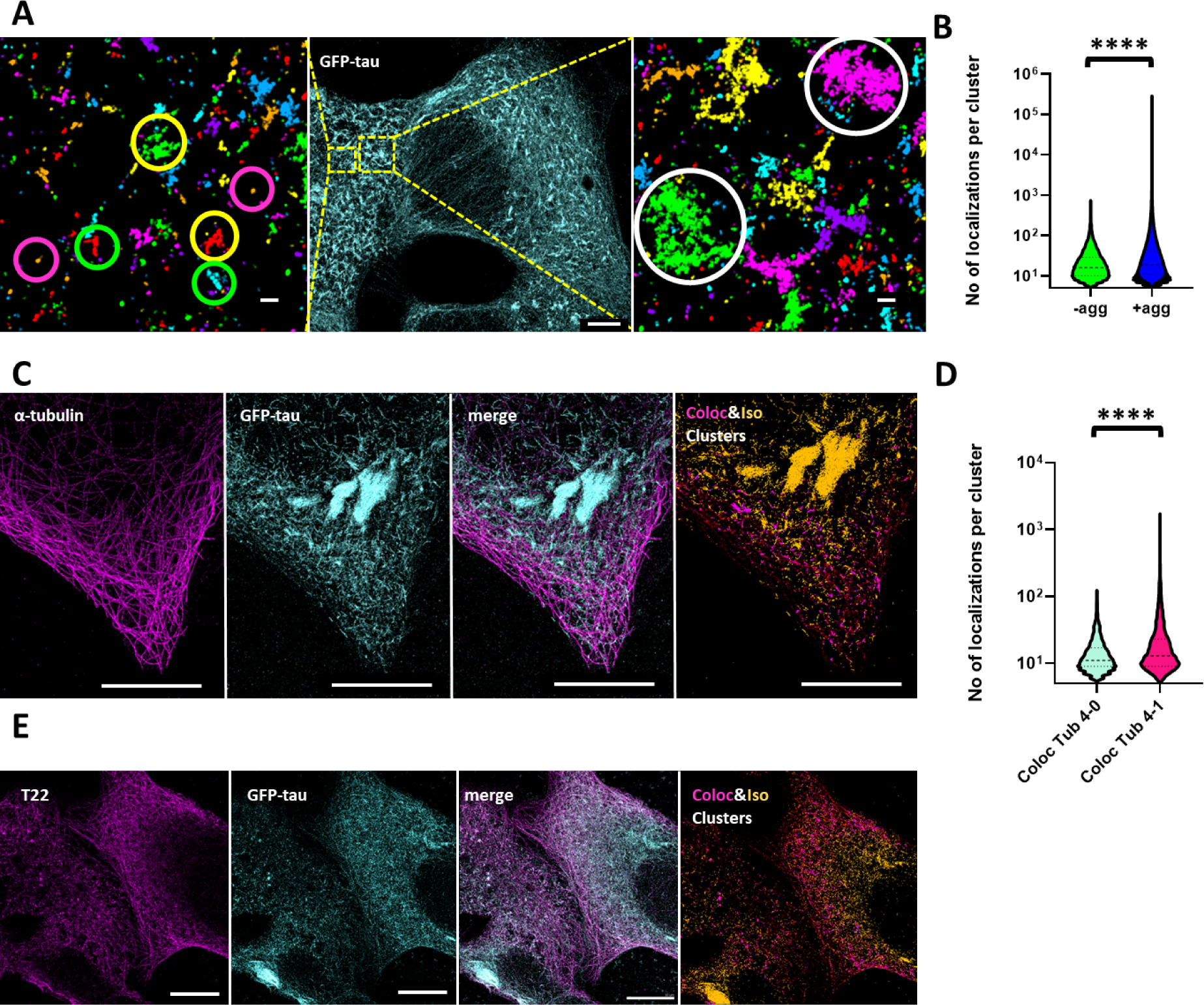
Pathological tau oligomers are distinct from physiological tau oligomers. **A**. Super-resolution image of tau in GFP-P301L tau (Clone 4.1) cells. Zoomed in regions after Voronoi segmentation are shown. Segmented images are pseudo color-coded with different colors corresponding to different segmented objects. Nano-clusters (magenta circles), fibrillary structures (green circles), branched fibrils (yellow circles) and conglomerate NFT-like structures (white circles) are visible in the segmented images. Scale bar in zoom out image is 5um and 200nm in zoom in images. **B**. Violin plots showing the number of localizations per Voronoi segmented tau nano-cluster in Clone 4.0 (-agg) and Voronoi segmented tau object in Clone 4.1 (+agg). The dashed lines indicate the median and the dotted lines indicate the 25^th^ and 75^th^ percentile (-agg: n=15 cells, n=3 experiments, +agg: n=20 cells, n=3 experiments). ****, *p*<0.0001. **C**. Two-color super-resolution images of α-tubulin (magenta), total tau (cyan) and overlay in GFP-P301L tau cells (Clone 4.1). The results of the co-localization analysis are shown in which tau aggregates co-localized with α-tubulin are color-coded in magenta and isolated aggregates are shown in yellow. Scale bars are 10um. **D**. Violin plots showing the number of localizations per Voronoi segmented tau nano-cluster that colocalizes with α-tubulin in Clones 4.0 and Clone 4.1. The dashed lines indicate the median and the dotted lines indicate the 25^th^ and 75^th^ percentile (Coloc Tub 4-0: n=9 cells, n=2 experiments, Coloc Tub 4-1: n=7 cells, n=2 experiments). ****, *p*<0.0001. **E**. Two-color super-resolution images of oligomeric tau detected by T22 (magenta), total tau (cyan) and overlay in GFP-P301L tau cells (Clone 4.1). The results of the co-localization analysis are shown in which tau aggregates co-localized with T22 are color-coded in magenta and isolated aggregates are shown in yellow. Scale bars are 10um.

To once again determine the percentage of oligomeric tau and the amount of cytosolic vs microtubule-associated tau present in these aggregated tau cells, we performed two-color SMLM. We labeled tau together with α-tubulin or with T22 and performed the co-localization analysis as for Clone 4.0. The percentage of microtubule-associated tau in Clone 4.1 (∼10±9% uncorrected or ∼17% corrected based on the co-localization positive control) was significantly lower than that in Clone 4.0 (∼61% uncorrected and 100% corrected) (Fig 1F). However, similar to Clone 4.0, ∼20±9% of tau (∼34% corrected) co-localized with the oligomeric T22 antibody (Fig 1F). The large tau aggregates reminiscent of neurofibrillary tangles were clearly excluded from microtubules and also did not co-localize with T22 (Fig 2C,E). The number of localizations per microtubule-associated tau object was higher in Clone 4.1 than that for the nano-clusters in Clone 4.0, suggesting that there are small tau aggregates that remain on the microtubule (Fig 2D). To further confirm this result, we next used the calibration approach to determine the copy number composition of the oligomeric tau species in Clone 4.1 by fitting the distribution of the number of localizations to the calibration function obtained from the nanobody dilution experiment (Fig S3D-G, Fig 2B and Fig S4C). The distribution of the number of localizations for Clone 4.1 is highly skewed with a long tail corresponding to the presence of a diverse range of large aggregates containing a high number of localizations. We therefore focused on fitting the part of the curve corresponding to smaller aggregates including nano-clusters and small fibrils. This analysis showed that the copy number of tau within these small aggregates ranged between 1-7 tau molecules. Interestingly, the percentage of monomeric tau was similar in Clone 4.0 and Clone 4.1 whereas the percentage of dimers and trimers decreased and the percentage of higher order oligomers increased in Clone 4.1 compared to Clone 4.0. These results suggest that the physiological dimers and trimers may seed the formation of the higher order pathological oligomers consisting of >3 tau proteins.

We next asked if tau aggregation had any impact on the integrity of the microtubule network. To address this question, we imaged the microtubule network in Clone 4.1 cells using SMLM. Surprisingly, tau aggregation led to disruption of the microtubule network leading to significantly less dense microtubules in Clone 4.1 compared to Clone 4.0 both with and without tau (Fig S1B-C, bottom panel and third plot). This finding suggests that tau aggregates have an adverse effect on the microtubule network integrity.

Overall, our results show the presence of oligomers consisting of >3 tau proteins in cells harboring tau aggregates that are not present under physiological conditions and that remain partially microtubule-associated, in addition to a diverse range of larger tau aggregates that together disrupt the integrity of the microtubule network in a cell model of FTDP-17.

### Iterative hierarchical clustering identifies the presence of distinct tau aggregate classes

To further investigate the types of higher order tau aggregates, present in Clone 4.1 cells, we developed an unsupervised shape classification algorithm based on iterative hierarchical clustering to classify the large tau aggregates present in SMLM images (see Methods) (Fig S5A-B). We named the algorithm, Iterative Hierarchical Clustering (constrained-relative standard deviation or c-RSD). We first visually separated Clone 4.1 cells into two categories: those showing low level of aggregation containing mainly small tau aggregates and those showing high level of aggregation containing larger tau aggregates. The SMLM images from these two categories were then segmented into individual tau aggregates using *density-based spatial clustering of applications with noise* (DBSCAN) ^28^ (Methods). We focused on tau aggregates containing more than 500 localizations as aggregates with fewer localizations represented a uniform class of nano-clusters with no particular shape (Fig S5C) (∼2.5% of all segmented objects had >500 localizations). DBSCAN segmentation of images from Clone 4.0 cells gave rise to very few segmented objects containing >500 localizations (∼0.12% of all segmented objects had >500 localizations), further confirming that these objects correspond to tau aggregates not present under physiological conditions. The tau aggregates coming from cells with low and high tau aggregation levels were combined into a single list and further classified with Iterative Hierarchical Clustering c-RSD. The number of localizations per tau aggregate, tau aggregate area, length and width were used as classification parameters. We imposed a threshold for the coefficient of variation for each parameter and grouped the tau aggregates together into individual classes as long as the coefficient of variation in the parameters did not exceed the imposed threshold (i.e. the tau aggregates within each class were “self-similar” within the imposed threshold) (Methods). Smaller variation thresholds result in a larger number of identified classes from the classification scheme. We selected the threshold such that the classification was more sensitive to aggregate length and width (i.e. aggregate shape) than the number of localizations and area.

Table 1 contains the number of tau aggregates that were segmented before classification and the resulting number of classes after classification. The classes were represented in the form of a dendrogram tree (Fig S5D). The initial classification tended to overestimate the number of classes and thus we further narrowed them down the by combining classes that were close in the dendrogram tree and visually looked similar to each other (Table 1, Fig S5E). This classification generated 22 classes containing tau aggregates coming from cells with both low and high levels of tau aggregation. These classes contained tau aggregates ranging in area by 3-orders of magnitudes (from 0.025 µm^2^ for the smallest classes to 30 µm^2^ for the largest ones). The presence of as many as 22 classes of tau aggregates suggests that the tau aggregates found in Clone 4.1 cells are highly diverse in terms of their size and shape. We plotted the results in the form of an ellipse graph, where each ellipse corresponds to a specific tau aggregate class (Fig 3A-J). The size of the ellipse corresponds to the percentage of total tau aggregates from cells with low or high levels of tau aggregation in that particular class (Table S1), whereas the shape of the ellipse represents the shape of the aggregates in that particular class (i.e. more elongated ellipses correspond to elongated aggregates with high aspect ratio). Visual inspection of aggregates within specific classes revealed linear tau fibrils (e.g. Class 6, Fig 3D), branched tau fibrils (e.g. Classes 9 and 10, Fig. 3E), NFT pre-tangle-like structures (e.g. Classes 12 and 15, Fig. 3I) as well as large agglomerate structures resembling NFTs (e.g Classes 21 and 22, Fig. 3J). These different classes may represent different stages of tau aggregation or different aggregation pathways. There was a large overlap in terms of the tau aggregate classes found in cells with low and high levels of tau aggregation. Linear and branched tau fibrils constituted the majority of tau aggregate classes in both cases. However, there were also some unique classes that were only present in cells with high levels of tau aggregation (Fig 3H,J, Table S1). In particular, classes having large areas and large number of localizations that resembled NFT-like structures (e.g. Classes 21 and 22) were only found in cells with high levels of tau aggregation whereas small tau fibrils (e.g. Class 6) were more representative of cells with low levels of tau aggregation.

**Table 1.** Number of tau aggregates and classes identified using Iterative Hierarchical Clustering c-RSD

| **Condition** | **Number of clusters**<br>**(before**<br>**classification)** | **Number of classes**<br>**(after unsupervised**<br>**classification)** | **Number of classes**<br>**(after manually**<br>**combining classes)** |
| --- | --- | --- | --- |
| Low tau aggregation | 1343 | N/A | N/A |
| High tau aggregation | 5539 | N/A | N/A |
| **Total** | **6882** | **40** | **22** |
| Thr231 staining | 225 | N/A | N/A |
| AT8 staining | 1913 | N/A | N/A |
| GFP nano staining | 6882 | N/A | N/A |
| **Total** | **9020** | **41** | **23** |

**Fig. 3.**
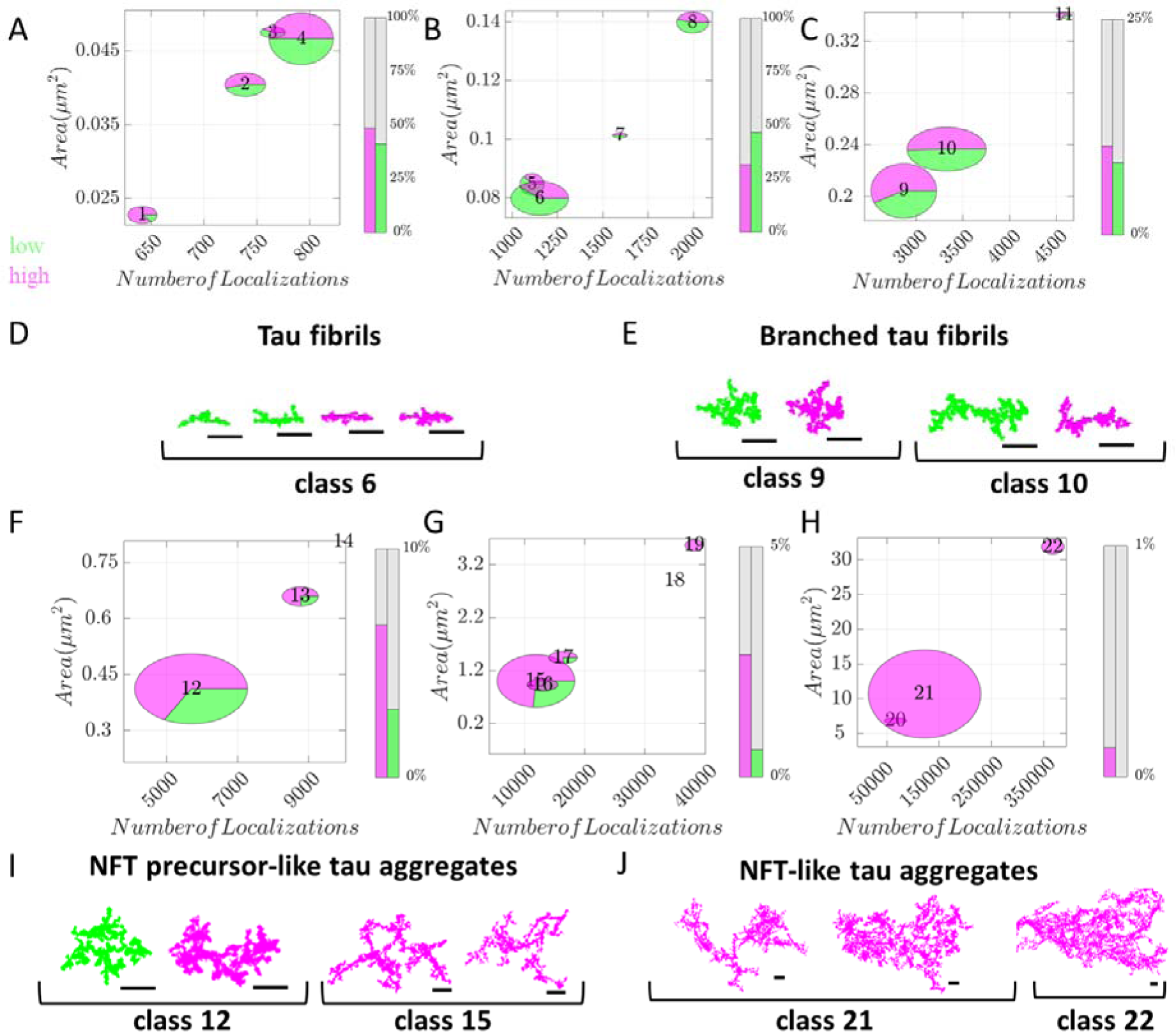
Shape classification reveals distinct classes of tau aggregates in Clone 4.1 cells. **A-C, F-H**. Ellipse plots showing the different tau classes found in low (green) and highly (magenta) aggregated tau cells. Each ellipse represents a separate class. The plot axes represent the number of localizations per tau aggregate and area of tau aggregates in µm^2^. Each ellipse is placed on the plot to represent the average number of localizations and area of the tau aggregates within that class. The size of the ellipses is scaled within each plot to represent the proportion of tau aggregates contained in that particular class but the size is not comparable between the different plots. Green represents the proportion of tau aggregates from low tau aggregation cells, whereas magenta represents the proportion of tau aggregates from high tau aggregation cells. The bar charts next to the ellipse plots represent the percentage of tau aggregates from each category (low in green and high in magenta) found in classes in the specified number of localizations and area range. Ellipses represent average numbers, not extreme outliers. **D, E, I, J**. Representative super-resolution images of tau aggregates from some of the most prominent classes in each plot. Shapes from low tau aggregation cells are colored green whereas those from high tau aggregation cells are colored magenta. Scale bars are 500nm.

### Phosphorylation of specific tau residues is associated with different types of higher order tau aggregates

Iterative Hierarchical Clustering c-RSD revealed the presence of a diverse range of tau aggregates (Fig 3A-J). However, the nature of tau present within these aggregates is unclear. In order to investigate the phosphorylation state of tau within these aggregate species, we performed two-color SMLM using two commonly used phosphor-tau antibodies, Thr231 and AT8, which are known to have high labeling specificity ^29^. Thr231 targets tau that is phosphorylated at the 231st threonine, which is considered to be an early marker of tau aggregation. AT8 targets tau that is phosphorylated at the serine residue at position 202 and threonine at position 205 (Ser202/Thr205), which is considered to be a late marker of tau aggregation. Staining with both antibodies gave rise to low, non-specific, fluorescent signal in Clone 4.0 cells compared to Clone 4.1 cells (Fig S6 A-B), indicating that tau present in Clone 4.0 cell line is lowly phosphorylated at any of the targeted residues ^30^. This result is in line with the expectation that hyper-phosphorylation of these residues is a marker of pathological tau aggregation and further supports that Clone 4.1 cells exhibit common markers of tau pathology found in disease. Single color and 2-color SMLM images of Clone 4.1 cells with either the Thr231 or the AT8 antibodies revealed that both antibodies bind to a wide range of tau aggregates having diverse shapes and sizes (Fig 4A-B, Fig S7A-B). Only ∼12±10% (∼20% corrected) of GFP-tau found within Clone 4.1 cells co-localized with either Thr231 or AT8 (Fig 1F), suggesting that aggregation in this cell model can potentially proceed in the absence of hyper-phosphorylation of these particular tau epitopes. The number of localizations per tau aggregate detected by the two antibodies was similar suggesting that there is overlap in the tau aggregates that these antibodies target (Fig S7A-C).

**Fig. 4.**
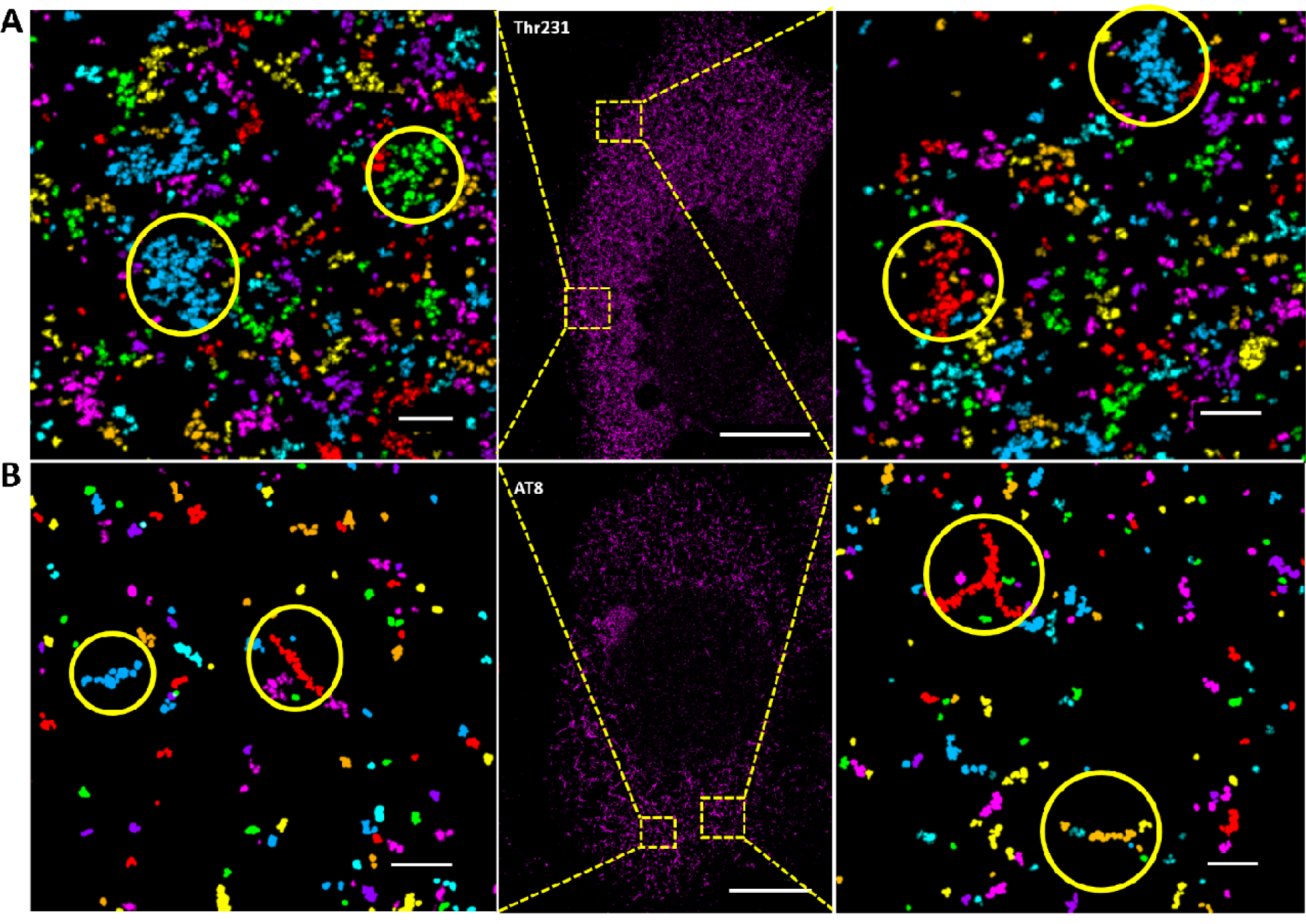
Branched tau fibrils and long tau fibrils are the predominant tau structure recognized by Thr231 and AT8, respectively. **A**. Super-resolution image of tau labelled with the Thr231 antibody in GFP-P301L tau (Clon 4.1) cells. Zoomed in regions after Voronoi segmentation are shown. Segmented images are pseudo color-coded with different colors corresponding to different segmented objects. Yellow circles highlight branched tau fibril like structures. Scale bar in zoom out image is 10um and 500nm in zoom in images. **B**. Super-resolution image of tau labeled with AT8 antibody in GFP-P301L tau (Clone 4.1) cells. Zoomed in regions after Voronoi segmentation are shown. Segmented images are pseudo color-coded with different colors corresponding to different segmented objects. Yellow circles highlight long tau fibril like structures. Scale bar in zoom out image is 10um and 500nm in zoom in images.

To further investigate whether different tau aggregates have differential phosphorylation states, we classified tau aggregates from Clone 4.1 cells stained with either the GFP nanobody or Thr231 or AT8 antibody using Iterative Hierarchical Clustering c-RSD. We obtained 23 classes, very similar to the number of classes obtained from classifying the GFP-nanobody labeled aggregates in cells having low or high levels of tau aggregation (Table 1). As expected, there was large overlap in the tau aggregate classes stained with Thr231, AT8 and GFP nanobody (Fig 5A-J). However, tau aggregates from cells stained with Thr231, which is an earlier tau aggregation marker, were over-represented by ∼2-fold in Class 3, corresponding to small branched tau fibrils (Table S2) compared to AT8 or GFP-nanobody. It is thus possible that phosphorylation at this residue is a characteristic of small branched tau fibrils. Similarly, AT8 labeled tau aggregates were over-represented by ∼2-4 fold in Classes 4 and 5, corresponding to long linear fibrillary structures, suggesting that phosphorylation at Ser202/Thr205 may be a prominent feature of this class of tau aggregates (Fig. 5D and Table S2). Furthermore, there were unique classes resembling NFT pre-tangle-like and NFT-like structures, which were only present in cells stained with the late tau aggregation marker AT8 but not the early tau aggregation marker Thr231 (Fig 5F-J, Table S2). It is possible that the phosphorylated Thr231 epitope is not accessible to the antibody within these large aggregates. However, it is also possible that not all tau aggregates acquire this phosphorylation mark.

**Fig. 5.**
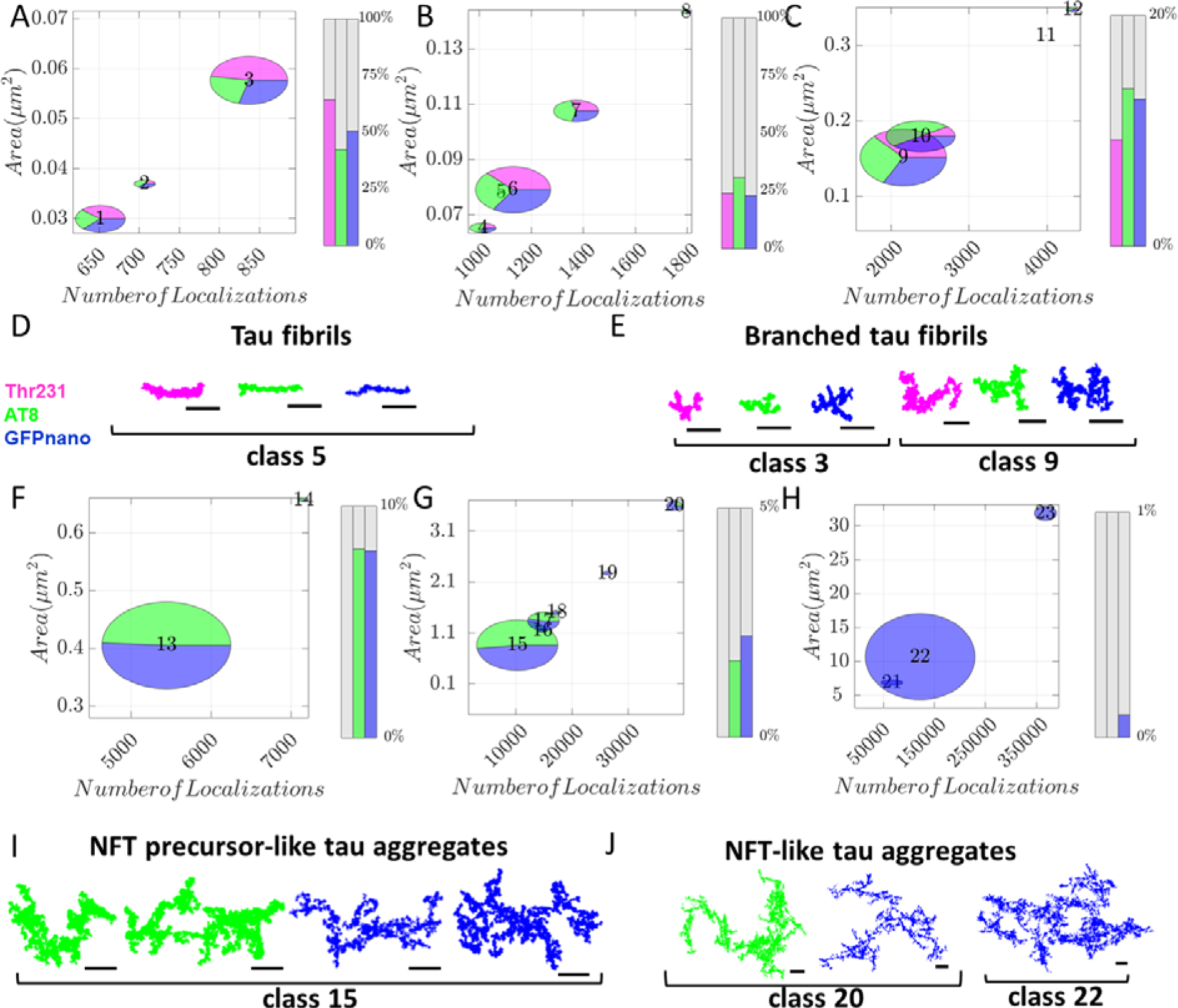
Shape classification differentiates higher order aggregates based on phosphorylation status. **A-C, F-H**. Ellipse plots showing the different tau classes found in Thr231 (magenta), AT8 (green) or GFP-nanobody (blue) labeled Clone 4.1 cells. Ellipse representation is the same as in Figure 3. **D, E, I, J**. Representative super-resolution images of tau aggregates from some of the most prominent classes in each plot. Shapes from Thr231 stained cells are colored magenta, those from AT8 stained cells are colored green and those from the GFP nanobody staining are colored blue. Scale bars are 500nm.

Overall, our results suggest the presence of overlapping as well as distinct classes of tau aggregates characterized by phosphorylation of different tau residues.

## Discussion

By using two cell models, one in which tau is predominantly microtubule associated and one that presents key features of disease pathology including the presence of aggregated tau inclusions and phospho-tau markers, we demonstrated using super-resolution microscopy, that tau forms physiological oligomers - mostly dimers and trimers - on microtubules, which are distinct from pathological oligomers and higher order aggregates. The distribution of tau on microtubules in the physiological cell models (WT-GFP-tau and P301L-GFP-tau in Clone 4.0) and in hippocampal neurons is not uniform, but instead we showed that tau forms nano-clusters.

Previous AFM and other *in vitro* studies have shown that P301L tau isolated from yeast^31^ and wild-type (WT) tau from rat^32^ form tau clusters and oligomers on taxol-stabilized microtubules, respectively. Further, these previous studies showed that P301L tau causes microtubules to stick together and bend abnormally, which was not the case for WT tau^31^. However, in the context of human tau expressed in mammalian cells and endogenous tau in neurons, we did not observe such microtubule abnormalities and both WT and P301L tau formed nano-clusters. Most recently, *in vitro* studies described the formation of physiological tau condensates^4^ also called tau “cohesive islands” ^5^ on microtubules. In both cases, these tau structures were shown to undergo liquid-liquid phase separation and the size of the condensates was in the micrometer range. The nano-clusters we observe are much smaller than these liquid condensates that form *in vitro*, instead, microtubule associated tau inside cells mainly consists of small tau complexes including monomers, dimers and trimers. Historically, tau oligomerization has been mostly deemed a pathological feature of tau. However, our findings show that tau can form small oligomers under physiological conditions. It has been speculated that an electrostatic zipper resulting from anti-parallel alignment of N-terminal halves of two tau molecules may be mediating tau dimerization^33^. Since tau is a highly disordered protein, it would be interesting to explore how it could form oligomers on microtubules in physiological conditions. Future studies exploring the relationship between different tau isoforms/mutants and physiological oligomer formation as well as investigating whether tau isoforms/mutants that don’t form physiological oligomers are resistant to aggregation would bring exciting new insights into the mechanisms of physiological and pathological tau oligomerization. From an evolutionary point of view, protein dimerization/oligomerization is a commonly used mechanism of functional regulation for many proteins as it promotes protein complex formation in physiological settings rather than just in pathology^34^. Our findings thus bring up interesting questions regarding the physiological function of tau oligomers, which should be subject of future studies. They can potentially mediate microtubule bundling as well as interact with and regulate the function of microtubule motors and severing enzymes as suggested in previous *in vitro* studies^35,4,5^.

Introducing tau oligomers into the brain of wild-type rodents has resulted in synaptic and memory dysfunction, supporting the hypothesis that tau oligomers are toxic^36,37^. Moreover, introducing tau oligomers directly into mammalian neurons using patch clamp electrodes also modified synaptic transmission and blocked events that could be underlying memory storage^38^. Interestingly, we showed that under pathological conditions the proportion of tau monomers was similar to physiological conditions but the proportion of dimers/trimers decreased and higher order oligomers appeared. This result demonstrates that there is a distinction between physiological and pathological tau oligomers. We further showed that physiological oligomers are not hyper-phosphorylated at residues Thr231 and Ser202 and Thr205 but pathological oligomers contain these phosphorylation markers, which reinforces the idea that physiological oligomers are distinct from pathological oligomers. Previous *in vitro* aggregation studies showed that tau dimers are the basic subunit of higher order tau oligomers^39^ and that tau dimerization is one important rate limiting step in the progression of tau aggregation^40^. These results are in line with our work, but here we further demonstrate that tau dimers/trimers exist in physiological conditions and thus may be primed to template further aggregation in disease.

We showed that the majority of pathological tau oligomers as well as higher order tau aggregates were excluded from microtubules. However, a small population of aggregated tau was found associated with microtubules, further suggesting that aggregation could be initiated from dimers/trimers present on microtubules. Hyperphosphorylation of tau has been considered a prerequisite for tau aggregation since the negative charge of phosphor residues neutralizes the positively charged microtubule-binding region of tau leading to tau’s dissociation from the microtubules and eventual aggregation^41^. However, tau oligomerization can occur without hyperphosphorylation or the addition of inducers suggesting that hyperphosphorylation of tau *in vivo* could be important for dissociation from microtubules but may not be necessary for the aggregation step^38^. Additionally, biochemical studies have shown that tau oligomers are most likely hetero-oligomers of non-hyperphosphorylated and hyperphosphorylated tau^42^. Hence, it is possible that tau aggregation can start on the microtubule before tau dissociation due to hyperphosphorylation. In the future, it would be interesting to study the kinetics of pathological tau aggregation over time to directly test this model. We further showed that tau aggregation negatively impacted the microtubule network itself. The density of the remaining microtubules in Clone 4.1 cells was lower than the basal conditions in Clone 4.0 cells (i.e. before the induction of tau aggregation). These results suggest that it is not just the dissociation of tau from the microtubule but the formation of tau aggregates that leads to microtubule disruption. It would be interesting to explore the mechanisms that lead to this microtubule network disruption and whether a specific tau aggregate species (oligomers or larger aggregates) leads to toxicity and loss of microtubule integrity.

Phosphorylation at Thr231 is an early aggregation event which diminishes the ability of tau to bind to microtubules *in situ* and is often used as a marker for postmortem diagnosis of most tauopathies^43^. Phosphorylation at Ser202 and Thr205 is a marker of late stage aggregation and is also used for postmortem diagnosis of tauopathies^44^. Here, we show that phosphor-tau antibodies that recognize these different phosphorylation sites label a subset of tau aggregates, suggesting that tau aggregation can proceed in the absence of phosphorylation at these residues. Further, using Iterative Hierarchical Clustering c-RSD, we classified the tau aggregates containing these specific phosphorylation marks. The resulting classification identified a variety of distinct classes of tau aggregates that could be grouped into 4 general categories: tau fibrils, branched tau fibrils, NFT pre-tangle-like tau aggregates and NFT-like tau aggregates. There was an overlap in the classes stained by the nanobody/antibodies but there were also classes that were more prominently recognized by either the Thr231 or the AT8 antibodies. These results suggest that phosphorylation at certain residues may be a characteristic of distinct types of tau aggregates. In recent years, the idea of tau strains with distinct structural conformations has become prevalent^45,46^. Amyloid β protein conformers have been characterized both *in vitro*^47^ and *in vivo*^48^ while some tau conformers have been characterized *in vitro* ^49,50^. However, the nature of these strains and which strains are responsible for which tauopathy or whether different strains lead to the formation of distinct tau aggregates are unknown. It is possible that some of the different classes of tau aggregates we detected in our FTDP-17 model are propagated by different tau strains, an idea reinforced by the finding that the phosphorylation state of tau is distinct in certain classes. It would be interesting in the future to determine whether different *MAPT* mutations associated with tauopathies lead to different tau aggregates.

Overall, we present a new approach based on super-resolution microscopy, quantitative analysis and unsupervised classification to characterize the distribution of tau in intact cells under physiological and pathological conditions with nanoscale spatial resolution. Our approach sheds new light on the physiological state of tau assemblies, which is poorly understood. It also opens the door for studying the mechanisms and kinetics of tau aggregation *in vivo*, the presence of early tau aggregates including pathological oligomers in disease and screening for drugs that can potentially target and disrupt these pathological oligomers. Further, our custom algorithm, Iterative Hierarchical Clustering c-RSD will be a useful tool for differentiating between tau aggregates associated with different tau strains in the future and could facilitate the study and identification of tau strains *in vitro* and *in vivo*.

## Conclusion

Here, using super-resolution microscopy, we show that tau forms oligomeric complexes on microtubules, which are distinct from pathological aggregates in a cell model of FTDP-17 as well as in neurons. Our findings suggest that microtubule-associated tau oligomers, specifically dimers and trimers, could template initial aggregation prior to dissociating from the microtubules. We thus show for the first time that super-resolution microscopy can detect and discriminate physiological and pathological tau oligomers, opening the door to study the early steps of tau aggregation *in vivo*. To further characterize the aggregation pathway in this FTDP-17 cell model, we developed an unsupervised shape classification algorithm, Iterative Hierarchical Clustering c-RSD for super-resolution microscopy images. We identified distinct classes of tau aggregates, characterized by early (Thr231) and late (Ser202/Thr205) tau aggregation markers as well as classes that lack either phosphorylation marker. This classification algorithm has the potential to differentiate between different tau aggregates formed by different tau strains in disease. Our work thus has important implications for studying mechanisms of tau aggregation in FTDP-17 and other tauopathies.

## Materials and Methods

### Cell culture

A stable cell line expressing WT-GFP 2N3R tau was derived from African green monkey (*Cercopithecus aethiops*) kidney epithelial cells (BS-C-1; CCL-26; American Type Culture Collection). Cells were grown in complete growth medium (Eagle’s minimum essential medium with Earle’s salts and non-essential amino acids plus 10% FBS, 1mM sodium pyriuvate, 2mM L-glutamine, 500 ug/ml Geneticin [G418 Sulfate; Thermo Fisher Scientific] and penicillin-streptomycin) in a 37°C incubator containing 5% CO_2_.

Two stable human embryonic kidney-derived QBI-293 cell lines (Clone 4.0, Clone 4.1) expressing full length human tau T40 (2N4R) carrying the P301L mutation with a GFP tag, a kind gift from the V. Lee lab at the University of Pennsylvania, were used for experiments. Clone 4.1 is sorted from Clone 4.0 cells to enrich for large compact tau aggregates after tau fibril addition. Cells were grown in Dulbecco’s modified Eagle’s medium supplemented with 10% tetracycline-screened fetal bovine serum (FBS), 1% pyruvate (10mM), 1% penicillin-streptomycin and L-glutamine (20 mM), 5ug/ml blasticidin, 200ug/ml Zeozin and were maintained in a 37°C incubator containing 5% CO_2_. Clone 4.1 was maintained in media containing 100ng/mL of doxycycline (Dox), whereas 100ng/nL of Dox was only added in the media 16 hours before fixation for Clone 4.0 in order to express the GFP-P301L tau construct. No tau fibrils were added in Clone 4.0 cells. For all imaging, cells were plated on eight-well Lab-Tek 1 coverglass chambers (Nunc).

E18 Sprague-Dawley rat hippocampal neurons were obtained in suspension from the Neuron Culture Service Center at the University of Pennsylvania. Neurons were plated in eight-well Lab-Tek 1 coverglass chambers (Nunc) which had been precoated with 0.5 mg/ml poly-L-lysine (Sigma-Aldrich) 24 hr before plating. Neurons were cultured in maintenance media consisting of Neurobasal (Gibco) supplemented with 2mM GlutaMAX, 100 U/ml penicillin, 100 mg/ml streptomycin, and 2% B27 (Thermo Fisher Scientific) in a 37°C incubator containing 5% CO_2_.

### Immunostaining

Two different fixation protocols were tested: paraformaldehyde (PFA) or methanol fixation. For PFA fixation, cells were initially washed with PBS for three times (5 min per wash), then fixed with 4% PFA for 10 min at room temperature and washed again three times (5 min per wash) with PBS following fixation. Since PFA fixation disrupted tau localization, for the experiments reported in the manuscript, cells were fixed using methanol fixation. Briefly, cells were first incubated with microtubule stabilizing buffer (MTSB: 15g PIPES, 1.9g EGTA, 1.32 MgSO_4_·7H_2_O, 5g KOH, H_2_O to a liter, pH=7) for 3 min and then ice cold methanol was added in the buffer for 3 min. Following this short incubation, cells were washed with MTSB twice. Cells were then blocked for 1hr using 4% (wt/vol) BSA in PBS. They were then incubated with the appropriate dilution of primary and secondary antibodies (or nanobody) in blocking buffer consisting of 3% (wt/vol) BSA and 0.2% Triton X-100 (vol/vol; Thermo Fisher Scientific) in PBS. Cells were washed with washing buffer (0.2% BSA and 0.05% Triton X-100; Thermo Fisher Scientific) between antibody/nanobody incubations. A list of antibodies/nanobodies used in this study is provided in Table S3. The nanobody we used was GFP VHH, recombinant binding protein (gt-250, Chromotek). The secondary antibodies we used were AffiniPure goat anti-mouse IgG (H+L, 115-005-003; Jackson ImmunoResearch) at a dilution of 1:100, and AffiniPure donkey anti-rabbit IgG (H+L, 711-005-152; Jackson ImmunoResearch) at a dilution of 1:100. The nanobody and the secondary antibodies used in this study were custom-labeled with an Alexa Fluor A647 or Alexa Fluor 405-Alexa Fluor 488 and an Alexa Fluor 405-Alexa Fluor A647 or Alexa Fluor 405-Alexa Fluor 488 activator/reporter dye pair combination at 0.12-0.15 mg/ml concentration, respectively ^51^.

### SMLM imaging

SMLM was acquired using an Oxford Nanoimaging system. The Nanoimager-S microscope had the following configuration: 405-, 488-, 561- and 640-nm lasers, 498-551- and 576-620-nm band-pass filters in channel 1, and 666-705-nm band-pass filters in channel 2, 100 × 1.4 NA oil immersion objective (Olympus), and a Hamamatsu Flash 4 V3 sCMOS camera. Localization microscopy images were acquired with 15 ms exposure for 50,000 frames. For multi-color SMLM, 110,000 frames with 15 ms exposure were acquired with sequential laser activation. The images were then processed using the NimOS localization software (Oxford Nanoimaging).

### Data analysis

#### Voronoi Tesselation Analysis

Localizations were exported in .csv format using the NimOS localization software and converted to .bin files using MATLAB R2017a. The rendered SMLM images were cropped using the AFIB plugin in ImageJ (National Institutes of Health) ^52^. Voronoi Tesselation Analysis was performed in MATLAB R2017a similarly to ^53^,^21^,^22^. First, a Voronoi threshold was chosen manually to define a Voronoi cluster as a collection of Voronoi polygons with areas smaller than the given threshold. We confirmed that the selected threshold led to proper segmentation of the super-resolution images visually and used the same threshold across different conditions for consistency. In the case of Clone 4.0, we cropped regions in which single microtubules were visible and avoided dense areas consisting of microtubule bundles to avoid segmentation errors due to inability to resolve the tau associated with individual microtubules within these bundles. The x and y coordinates from the localizations were processed by the “delaunayTriangulation” function and then the ‘Voronoidiagram” function to generate Voronoi polygons. The Voronoi polygon areas were calculated from the shoelace algorithm. A new molecule list assigning localizations to different channels (0-9) according to their Voronoi area was then generated. Next, these localizations were put into different clusters and their cluster statistics were generated and used for analysis. The different clusters were pseudocolored based on their channel (0-9) and visualized using a custom-written software, Insight3 (provided by B. Huang, University of California, San Francisco, San Francisco, CA;^54^.

#### Outlier Removal Analysis

In the segmented clusters, there is a positive correlation between the cluster area and the number of localizations per cluster. However, clusters corresponding to imaging artefacts (e.g. molecules that do not photoswitch) fall outside of this positive correlation as they contain a large number of localizations but small area. We used this positive correlation between cluster area and the number of localizations to filter clusters corresponding to imaging artefacts. This procedure removed 0.1316% of clusters from the data.

#### GFP nanobody Calibration

We determined the number of localizations corresponding to a single tau molecule labeled with the GFP-nanobody by carrying out super-resolution imaging under dilute labeling conditions corresponding to a 1:10000 dilution of the GFP-nanobody. We segmented these images using Voronoi segmentation as described above and determined the number of localizations per cluster. The number of localizations distribution was fit to a log normal function to extract two parameters mu and sigma corresponding to the log normal distribution function (f1). The above-mentioned distribution function (f1) and its convolution with itself two times (f2 = f1*f1) and three times (f3 = f2*f1) is used to determine the percentage of single-tau molecules, monomers and oligomers by fitting a linear combination of these functions to the distribution of the number of localizations per cluster from the actual experiment corresponding to 1:100 dilution of the GFP-nanobody as previously described ^25^.

#### SMLM Colocalization Analysis

Colocalization analysis was performed using a custom-built algorithm in MATLAB R2017a for all multi-color SMLM data. Regions with similar tau density were cropped for quantitative analysis, avoiding overcrowded regions to minimize segmentation errors. The threshold on Voronoi polygon size was kept consistent between different conditions and clusters having fewer than 5 localizations were filtered. First, a list of reference boundaries from the reference cluster (channel 1) is obtained. The boundaries are then extended by a set radius (30 nm) and the points inside the boundary are obtained. Next, the points from the non-reference cluster (channel 2) that fall into the reference cluster are determined. The percentage of colocalized points in the non-reference clusters is also determined. A threshold of 40% colocalization is required for a cluster to be considered colocalized and anything below 40% is considered isolated. This threshold was determined after qualitative examination of the colocalized clusters visually and used throughout the analysis for consistency.

#### Distance Distribution Correction (DDC) Analysis

The DDC algorithm developed by ^24^ was used to correct for blinking in SMLM images acquired using the GFP nanobody fused to Alexa Fluor A647 or Alexa Fluor 405-Alexa Fluor 488 without 405nm activation as previously explained.

#### Iterative Hierarchical Clustering (constrained-relative standard deviation (c-RSD)

In order to segment the localized points in the super-resolution images into “objects” we performed a density-based clustering algorithm (DBSCAN) ^28^. DBSCAN requires two input parameters for segmentation: the minimum number of points (k) within a search radius (epsilon). We chose a minimum number of five localizations per object as a threshold for the segmentation (k = 5). To determine the search radius (epsilon) in an unbiased manner, we employed the elbow method. We first calculated the 4^th^ nearest neighbor distances (NNDs) (k-1) between each of the localizations in the entire STORM image. We then sorted and plotted the NNDs and determined the value corresponding to the elbow point of this sorted distribution. The distance corresponding to the elbow point was used as the search radius in the DBSCAN algorithm. After running the DBSCAN algorithm on the entire STORM image, localizations were segmented into objects. In order to classify these objects into unique classes we next extracted their features. In our classification algorithm we considered 8 features describing each object. To determine the most relevant features to use in the classification, we first performed a PCA analysis on these objects to obtain the two axes corresponding to the most variation between the localization points. The 8 features were the number of localizations per object, object area, number of localizations in the upper-left, upper-right, lower-left, and lower-right quarters of the object and the length corresponding to the major and minor axes of the object. To classify the objects based on these 8 features, we used agglomerative Hierarchical clustering algorithm ^55^. In Hierarchical clustering, there is no a priori assumption on the number of classes. We found that the Hierarchical clustering alone over-estimated the number of classes, and often objects having very similar features were classified into distinct classes. In order to overcome this problem, we developed an iterative Hierarchical clustering algorithm, that we named Iterative Hierarchical Clustering (constrained-relative standard deviation (c-RSD)). We imposed a threshold on the coefficient of variation (ratio of the standard deviation to the mean) on two of the features (length and width of the object). First, we constructed a linkage (dendrogram tree) by calculating the distance between each of the objects in the 8^th^ dimensional space. After forming the dendrogram tree, if we have N objects to classify, the closest two objects and the remaining (N-2)-many closest objects from these two first closest objects can be obtained. As a result, we can write a set wherein the N-many objects are sorted based on their distances from the first two closest objects. After doing so the algorithm combines the k-closest objects into a single class if and only if the coefficient of variation on the major and minor axis length of the new class is still less than or equal to the imposed threshold on the coefficient of variation (0.15). In the next step, we will perform the above-mentioned algorithm iteratively on the remaining classes until the number of classes reaches a plateau. It should be mentioned that in the second loop and so on, the dendrogram tree is formed from the average value of the features in a class and the restriction on the coefficient of variation is always set on the objects within a class.

As a final step we can furthermore classify the classes obtained from the Iterative Hierarchical Clustering c-RSD into unique classes based on the visual inspection of these classes. Any classes that visually resembled each other and that were connected in the dendrogram tree were furthermore combined together. Calculation of the coefficient of variation on the major/minor axis length after this final classification step might be higher than the 0.15 value because of the supervised step but visually they appear to belong to the same class.

### Statistical Analysis

Non-parametric Mann-Whitney tests were performed using OriginPro 2017 software for all data imaging data sets. A paired student t-test was performed for the fixation control experiment in Fig. S2B.

## Acknowledgments

We thank Prof. V. Lee, UPenn for the QBI-293 cell lines (Clone 4.0 and 4.1) expressing full length human tau T40 (2N4R) carrying the P301L mutation with a GFP tag. We thank Dr. P. K. Relich for writing the co-localization analysis code while at UPenn. We thank Dr. Angel Sandoval Alvarez, ICFO, Barcelona, for making the cell lines expressing wild type tau fused to GFP. We thank Dr. C. Evans from the Holzbaur lab, UPenn for her help with culturing neurons. We thank Dr. E. Lee, Dr. L. Rhoades and Dr. P. K. Relich for their valuable input on the manuscript. This work was funded by a Linda Pechenik Montague Investigator Award, Penn Institute on Aging Pilot Grant Award and NIH/NIGMS R01GM133842 to M.L.

## Author Contributions

M.L. and M.T.G. conceived of the study and designed experiments. M.T.G. carried out the experiments and analyzed the data. A.A. developed and wrote the Iterative Hierarchical Clustering (c-RSD) code. E.M.S. helped with sample preparation including custom fluorescence labeling of nanobody. A.N.S.R. helped with multi-color experiments using Thr232 and AT8 antibodies. C.H.B wrote the code for DDC correction and analyzed data with DDC. J.X. supervised C.H.B with the development of DDC correction algorithm. M.L. supervised the study and acquired funding. M.T.G. and M.L. wrote the first draft of the manuscript. All authors provided feedback on the final manuscript.

