## Supplementary Materials for "Tau forms oligomeric complexes on microtubules that are distinct from pathological oligomers in disease"

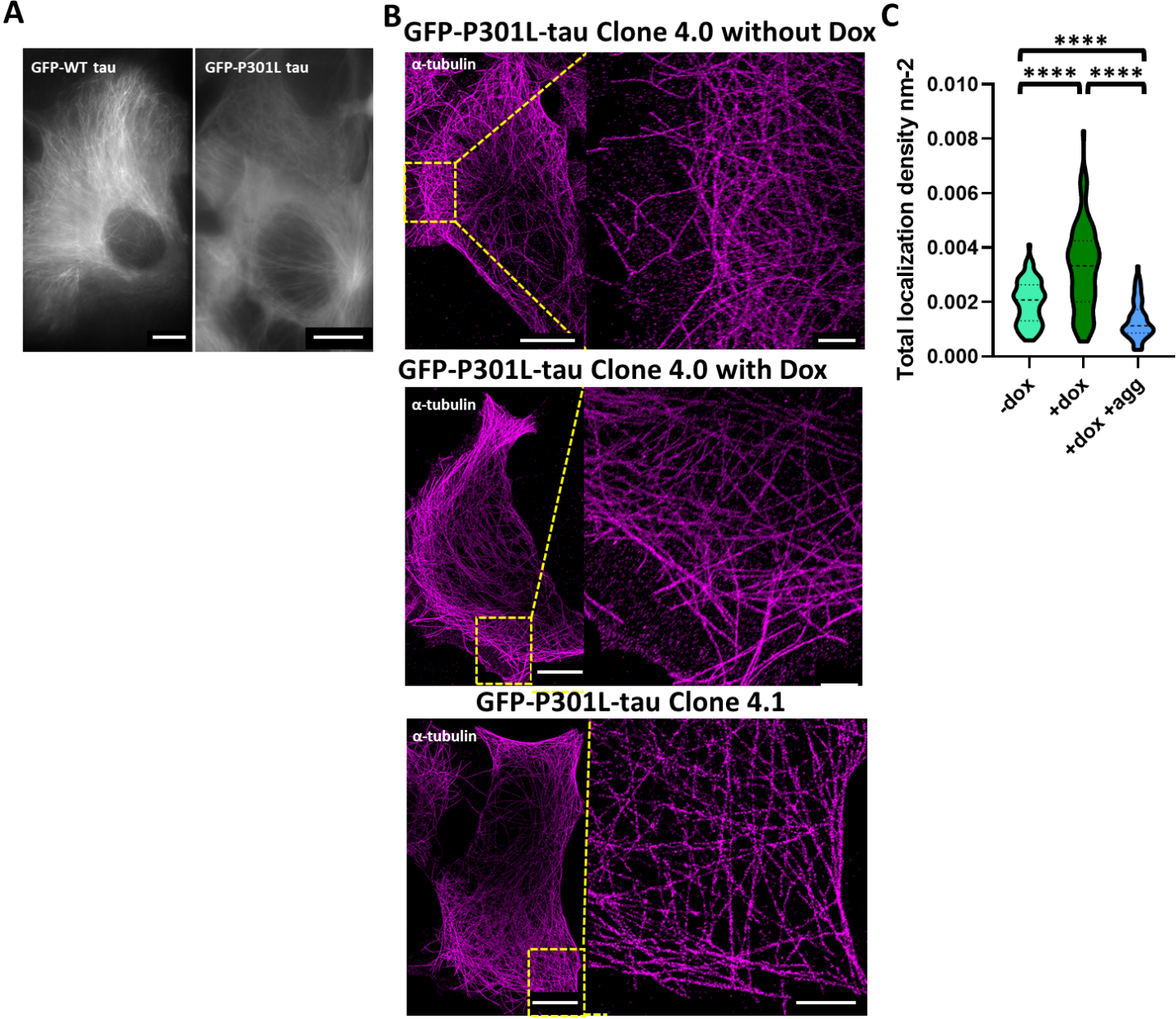

**Fig. S1. The microtubule network in GFP-WT tau and GFP-P301L tau (Clone 4.0 and 4.1) cells**

**A.** Wide-field images of GFP fluorescence in live cells expressing GFP-WT tau and GFP-P301L tau (Clone 4.0) cells. Scale bars are 10 um.

**B.** Super-resolution image and zoom of α-tubulin in Clone 4.0 cells in the absence of Dox induction (i.e no GFP-tau expression) (top panel). Super-resolution image and zoom of α-tubulin in Clone 4.0 cells in the presence of Dox induction (i.e non-aggregated tau expression) (middle panel). Super-resolution image and zoom of α-tubulin in Clone 4.1 cells in the presence of Dox induction (i.e aggregated tau expression) (bottom panel). Scale bar for zoom out images is 10um and 2um for zoom in images.

**C.** Violin plots showing the total tubulin localization density per nm^2^ in the three different conditions in A-C. Tubulin localization density is proportional to microtubule network density. The dashed lines indicate the median and the dotted lines indicate the 25^th^ and 75^th^ percentile (-dox: n=12 cells, n=2 experiments, +dox: n=10 cells, n=2 experiments, +dox +agg: n=9 cells, n=2 experiments). ****, *p*<0.0001.

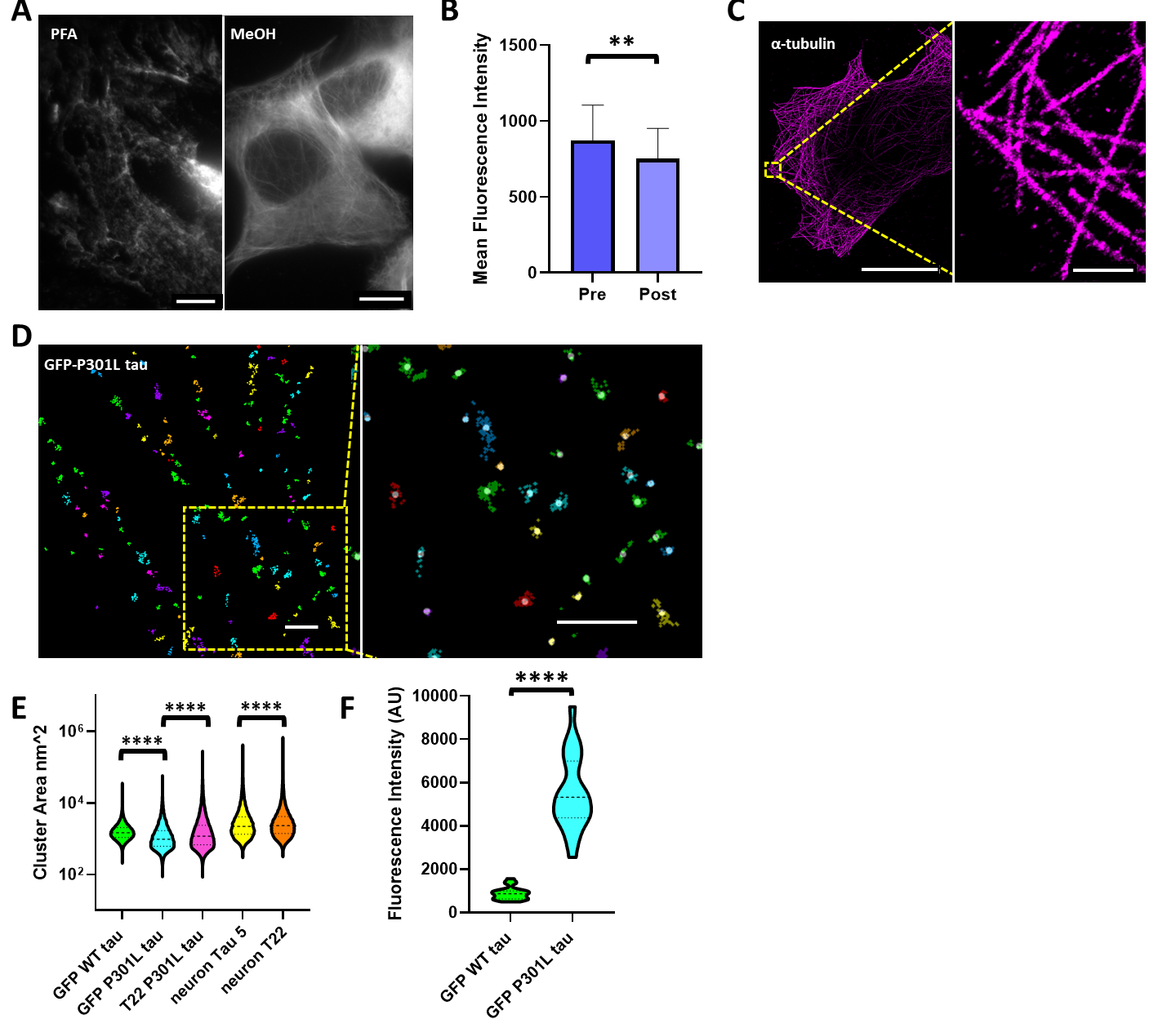

**Fig. S2. Tau forms nano-clusters on microtubules in a number of different cell models and in hippocampal neurons**

**A.** Wide-field images of GFP fluorescence in cells expressing GFP-P301L tau (Clone 4.0) fixed with either PFA or MeOH. Scale bars are 10 um.

**B.** Bar plots showing the mean fluorescence intensity of GFP-tau in AU before (pre-) and (post-) methanol fixation in GFP-P301L-tau cells (Clone 4.0) (n=5 cells). **, *p*<0.01.

**C.** Super-resolution image of α-tubulin staining in GFP-P301L-tau (Clone 4.0) cells and zoom. Scale bar in zoom out image is 20um and 1um in zoom in image.

**D.** Voronoi segmentation of a region of a GFP-P301L-tau (Clone 4.0) cell and zoom. The different colors represent different nano-clusters whose center is represented by a white dot. Individual fluorophore localizations are represented as color-coded crosses. Localizations that are in close spatial proximity are segmented as belonging to the same nano-cluster. Scale bars are 200nm in both zoom out and zoom in images.

**E.** Violin plots showing the area of Voronoi segmented nano-clusters in nm^2^ in the different cell lines used in this study. The dashed lines indicate the median and the dotted lines indicate the 25^th^ and 75^th^ percentile (GFP-WT tau: n=15 cells, n=2 experiments, GFP-P301L tau: n=15 cells, n= 3 experiments, T22 P301L tau: n=19 cells, n=3 experiments, neuron Tau 5: n=3 cells, neuron T22: n=3 cells). ****, *p*<0.0001.

**F.** Violin plots showing the mean fluorescence intensity (AU) in conventional fluorescence images of tau in GFP-WT-tau and GFP-P301L-tau live cells. The fluorescence intensity of GFP-tau is proportional to the tau expression level in the two cell lines. The dashed lines indicate the median and the dotted lines indicate the 25^th^ and 75^th^ percentile (GFP-WT tau: n=20 cells, GFP-P301L tau: n=20 cells). ****, *p*<0.0001.

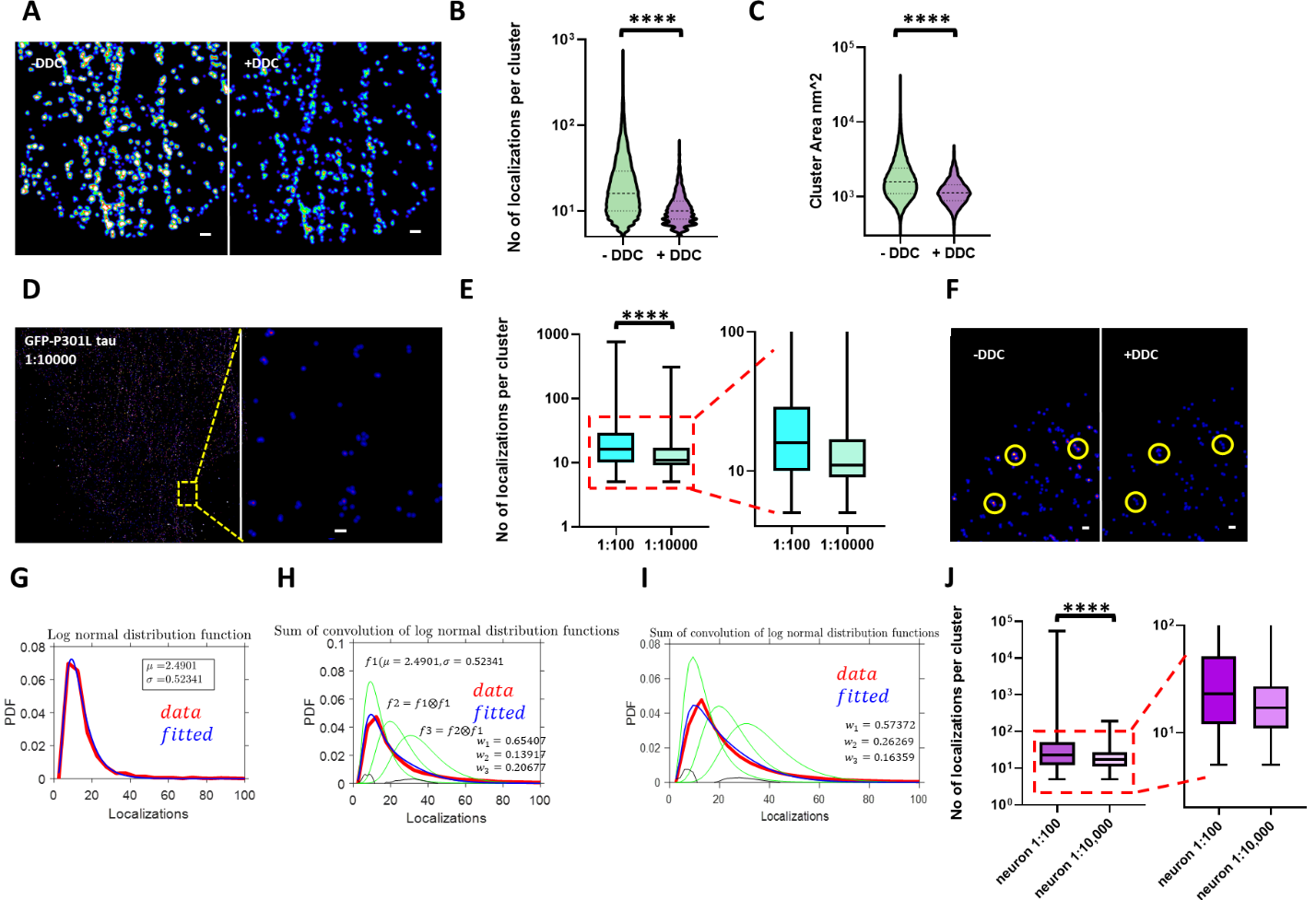

**Fig. S3. Tau nano-clusters present in the GFP-P301L-tau cell line (Clone 4.0) mostly consists of monomers, dimers and trimers**

**A.** Super-resolution images of GFP-tau in Clone 4.0 before and after DDC correction. Color map corresponds to localization density with less dense regions shown in blue and denser regions shown in red. Scale bars are 100nm in both images.

**B.** Violin plots showing the number of localizations per Voronoi segmented nano-cluster in Clone 4.0 before and after DDC correction. The dashed lines indicate the median and the dotted lines indicate the 25^th^ and 75^th^ percentile (n=15 cells, n=3 experiments). ****, *p*<0.0001.

**C.** Violin plots showing the area of Voronoi segmented nano-clusters in nm^2^ in Clone 4.0 before and after DDC correction. The dashed lines indicate the median and the dotted lines indicate the 25^th^ and 75^th^ percentile (n=15 cells, n=3 experiments). ****, *p*<0.0001.

**D.** Super-resolution image of GFP-P301L tau in Clone 4.0, where the GFP nanobody has been diluted 100-fold more than in the usual imaging conditions and zoom. Color map corresponds to localization density with less dense regions shown in blue and denser regions shown in red. Scale bar is 100nm.

**E.** Box plots showing the number of localizations per Voronoi segmented nano-cluster in normal labeling conditions corresponding to 1:100 dilution of the GFP nanobody and in dilute labeling conditions corresponding to 1:10,000 dilution of the nanobody. (1:100: n=15 cells, n=3 experiments, 1:10,000: n=5 cells). Box corresponds to 25-75 percentile, line corresponds to median and whiskers correspond to the minimum and maximum. A zoom of the box plot is shown as inset. ****, *p*<0.0001.

**F.** Super-resolution images of GFP-P301L tau in Clone 4.0, where the GFP nanobody has been diluted 100-fold more than in the usual imaging conditions, before and after DDC correction. Color map corresponds to localization density with less dense regions shown in blue and denser regions shown in red. Yellow circles indicate GFP tau nano-clusters before and after DDC correction. Scale bars are 100nm in both images.

**G.** Plot showing the number of localizations per nano-cluster under dilute labeling conditions (red) and the log normal fit (blue) used as calibration function (f_1_) for monomeric tau.

**H.** Plot showing the number of localizations per nano-cluster under normal (experimental) labeling conditions (red) in Clone 4.0 cells (GFP-P301L), dimeric and trimeric calibration functions (f_2_ and f_3_) (green) obtained by linear convolution of the monomeric calibration function (f_1_) obtained from the log normal fit in (G), and the fit of the experimental data to a combination of f_1_, f_2_ and f_3_ (blue) with weights w_1_, w_2_ and w_3_ corresponding to proportion of monomers, dimers and trimers.

**I.** Plot showing the number of localizations per nano-cluster under normal (experimental) labeling conditions (red) in GFP-WT cells, dimeric and trimeric calibration functions (f_2_ and f_3_) (green) obtained by linear convolution of the monomeric calibration function (f_1_) obtained from the log normal fit in (G), and the fit of the experimental data to a combination of f_1_, f_2_ and f_3_ (blue) with weights w_1_, w_2_ and w_3_ corresponding to proportion of monomers, dimers and trimers.

**J.** Box plots showing the number of localizations per Voronoi segmented nano-cluster in normal labeling conditions corresponding to 1:100 dilution of the Tau 5 primary antibody and the secondary anti-mouse antibody labeled with Alexa 647 and in dilute labeling conditions corresponding to 1:10,000 dilution of the Tau 5 primary antibody and 1:100 dilution of the anti-mouse secondary antibody labeled with Alexa 647. (1:100: n=7 cells, n=2 experiments, 1:10,000: n=6 cells, n=2 experiments). Box corresponds to 25-75 percentile, line corresponds to median and whiskers correspond to the minimum and maximum. A zoom of the box plot is shown as inset. ****, *p*<0.0001.

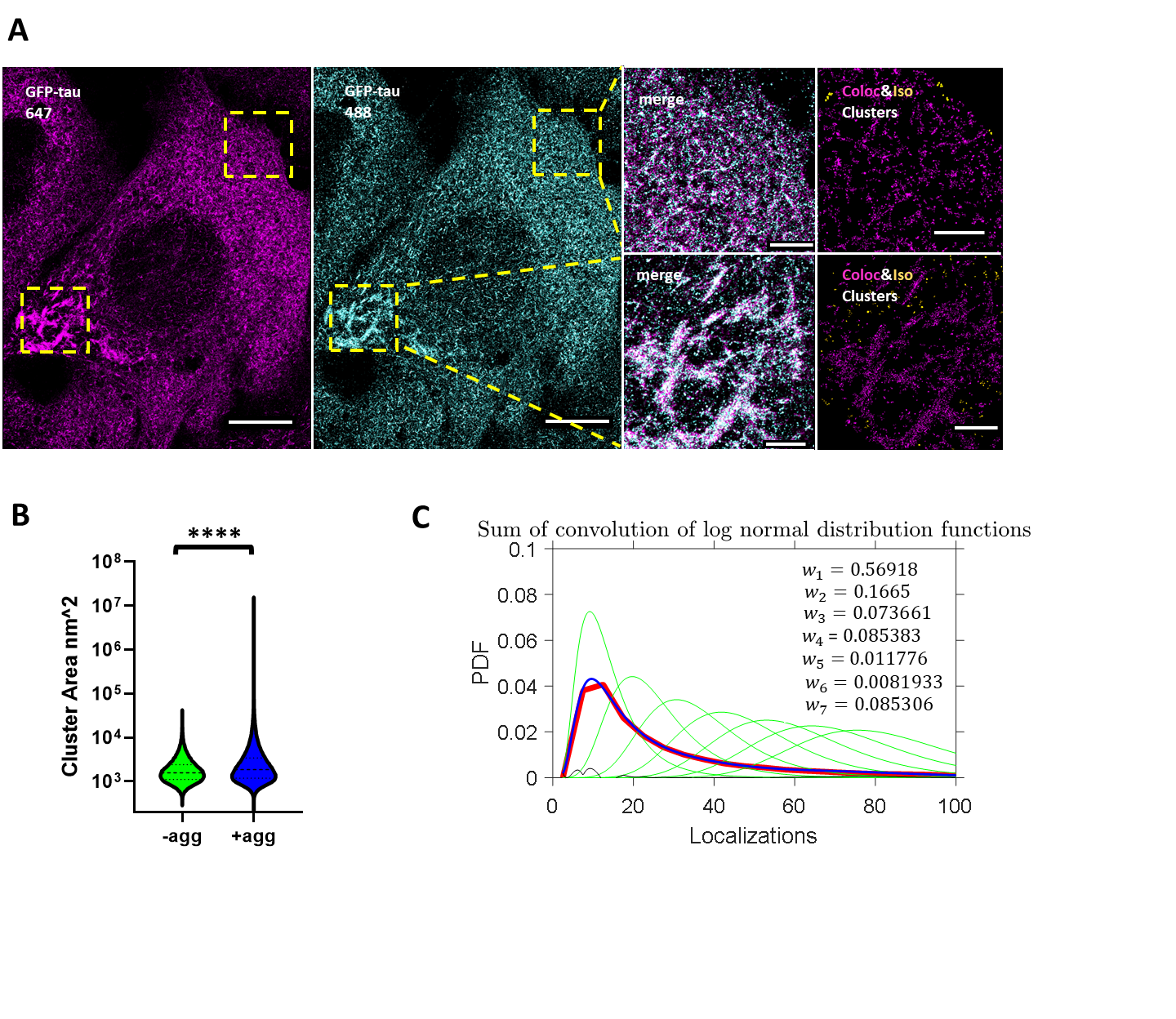

**Fig. S4. Pathological tau oligomers are larger in size than physiological tau oligomers**

**A.** Two-color super-resolution images of GFP-tau stained with GFP nanobody fused to Alexa-647 fluorophore (magenta), or Alexa 488 fluorophore (cyan) and zoomed in overlay of two different regions (yellow boxes) in GFP-P301L tau cells (Clone 4.1). The results of the co-localization analysis are shown in which tau nano-clusters (488) co-localized with GFP-tau (647) are color-coded in magenta and isolated tau nano-clusters are shown in yellow. Scale bars are 10 um in the first two images and 2um in the zoomed in images.

**B.** Violin plots showing the area of Voronoi segmented nano-clusters in nm^2^ in Clones 4.0 (-agg) and Voronoi segmented tau objects in Clone 4.1 (+agg). The dashed lines indicate the median and the dotted lines indicate the 25^th^ and 75^th^ percentile (-agg: n=15 cells, n=3 experiments, +agg: n=20 cells, n=3 experiments). ****, *p*<0.0001.

**C.** Plot showing the number of localizations per tau object under normal (experimental) labeling conditions (red) in Clone 4.1 cells (GFP-P301L), dimeric-heptameric calibration functions (f_2_-f_7_) (green) obtained by linear convolution of the monomeric calibration function (f_1_) obtained from the log normal fit in (Fig. S2F), and the fit of the experimental data to a combination of f_1_-f_7_ (blue) with weights w_1_-w_7_ corresponding to proportion of monomers, dimers, trimers and higher order oligomers.

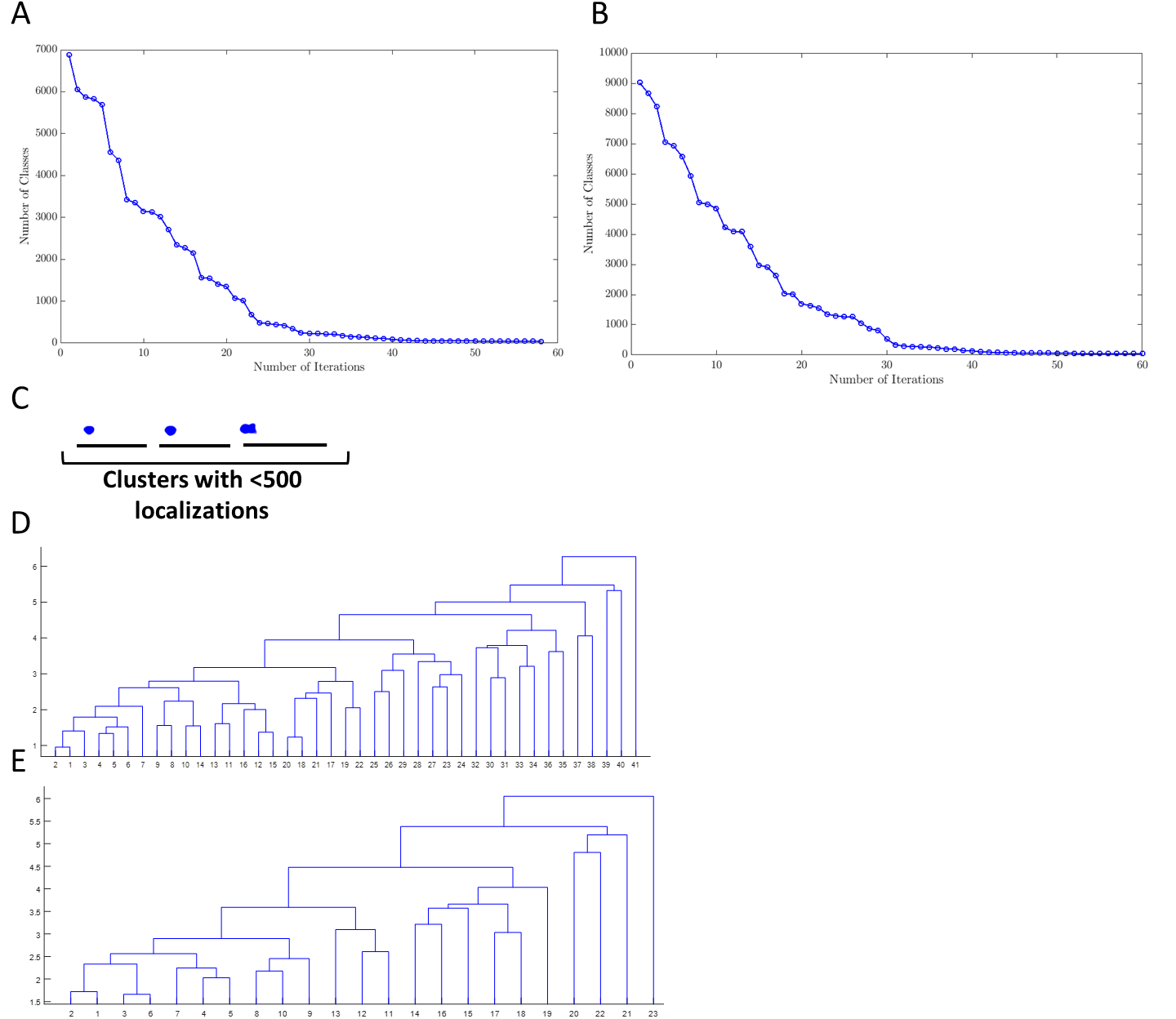

**Fig. S5. Iterative hierarchical shape classification**

**A.** Plots showing the number of classes identified after each iteration of the hierarchical clustering algorithm in low and high tau aggregation cells. The number of classes identified converge after about 30 iterations.

**B.** Plots showing the number of classes identified after each iteration of the hierarchical clustering algorithm in GFP-nanobody, Th231 and AT8 labeled cells. The number of classes identified converge after about 30 iterations.

**C.** Examples of clusters having <500 localizations in aggregated (Clone 4.1) cells. These structures mostly represent nano-clusters with no particular shape. Scale bars are 500nm.

**D.** Dendrogram tree resulting from unsupervised classification showing 41 classes of tau aggregates and how they relate to each other. This example is derived from the analysis of Thr231, AT8 and GFP nano stained Clone 4.1 cells. The x axis represents the number of classes, whereas the y axis represents the z-score/height.

**E.** Dendrogram tree after manually combining the 41 classes resulting from the unsupervised classification into 23 classes of tau aggregates and how they relate to each other. This example is derived from the analysis of Thr231, AT8 and GFP nano stained Clone 4.1 cells. The x axis represents the number of classes, whereas the y axis represents the z-score/height.

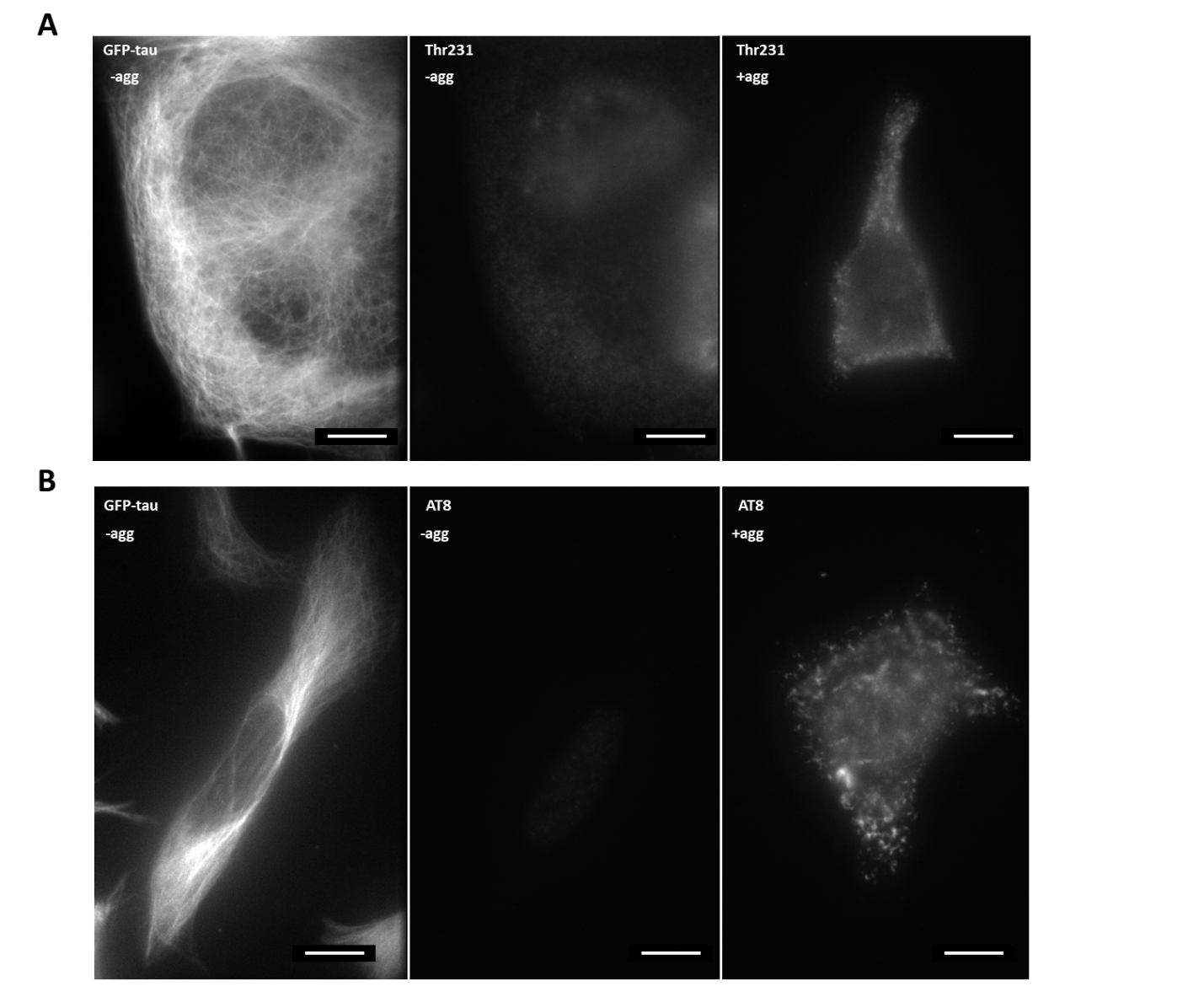

**Fig. S6. Thr231 and AT8 antibodies specifically stain tau in Clone 4.1 but not Clone 4.0**

**A.** Wide-field images of GFP fluorescence in Clone 4.0 (GFP-tau –agg, left panel) and fluorescence signal from Thr231 antibody staining (Thr231 –agg, middle panel) in the same cell. Fluorescence signal from Thr231 antibody staining in a Clone 4.1 cell (Thr231 +agg, right panel). Scale bars are 10um.

**B.** Wide-field images of GFP fluorescence in Clone 4.0 (GFP-tau –agg, left panel) and fluorescence signal from AT8 antibody staining (AT8 –agg, middle panel) in the same cell. Fluorescence signal from AT8 antibody staining in a Clone 4.1 cell (AT8 +agg, right panel). Scale bars are 10um.

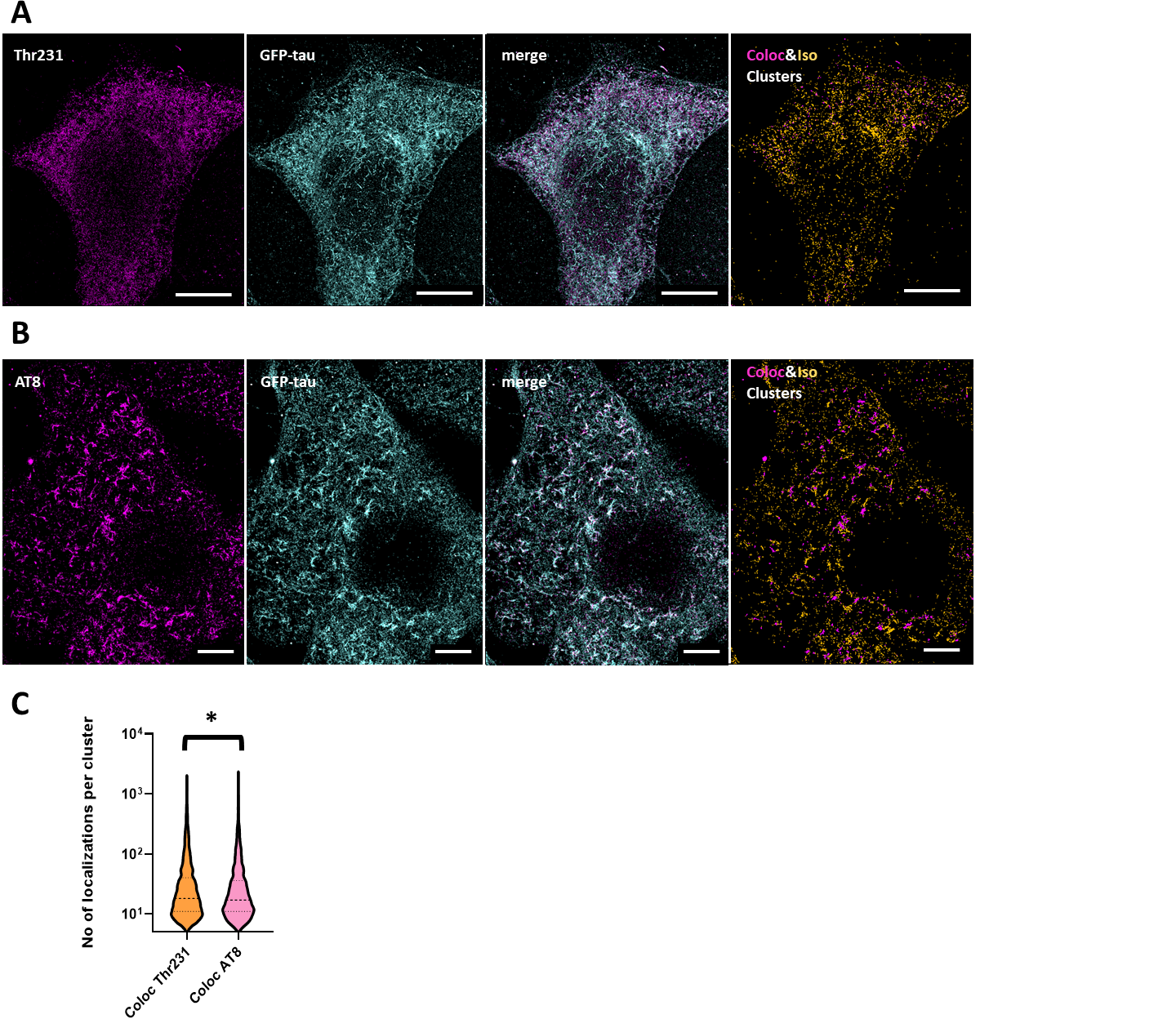

**Fig. S7. Phosphorylation of specific tau residues is associated with a diverse range of tau aggregates**

**A.** Two-color super-resolution images of Thr231 (magenta), total tau (cyan) and overlay in GFP-P301L tau cells (Clone 4.1). The results of the co-localization analysis are shown in which tau co-localized with Thr231 is shown in magenta and isolated tau is shown in yellow. Scale bars are 10um.

**B.** Two-color super-resolution images of Thr231 (magenta), total tau (cyan) and overlay in GFP-P301L tau cells (Clone 4.1). The results of the co-localization analysis are shown in which tau co-localized with Thr231 is shown in magenta and isolated tau is shown in yellow. Scale bars are 5um.

**C.** Violin plots showing the number of localizations per segmented Voronoi object of GFP-tau that colocalizes with Thr231 or AT8 in Clone 4.1. The dashed lines indicate the median and the dotted lines indicate the 25^th^ and 75^th^ percentile (Coloc Thr231: n=6 cells, n=2 experiments, Coloc AT8: n=5 cells, n=2 experiments). *, *p*<0.05.

| **Class** | **Number of clusters per class** | **Ratio of low tau aggregation clusters over the total no of low tau aggregation clusters in each class** | **Ratio of high tau aggregation clusters over the total no of high tau aggregation clusters in each class** |
| --- | --- | --- | --- |
| 1 | 598 | 0.01787044 | 0.10362881 |
| 2 | 827 | 0.10275503 | 0.12439068 |
| 3 | 507 | 0.10052122 | 0.06716014 |
| 4 | 1311 | 0.19136262 | 0.19028706 |
| 5 | 446 | 0.09828742 | 0.05668893 |
| 6 | 1066 | 0.2278481 | 0.13720888 |
| 7 | 276 | 0.04095309 | 0.0398989 |
| 8 | 595 | 0.10052122 | 0.08304748 |
| 9 | 281 | 0.03201787 | 0.04296804 |
| 10 | 337 | 0.0476545 | 0.04928687 |
| 11 | 67 | 0.00446761 | 0.01101282 |
| 12 | 302 | 0.02382725 | 0.04874526 |
| 13 | 97 | 0.00521221 | 0.01624842 |
| 14 | 10 | 0.0007446 | 0.00162484 |
| 15 | 76 | 0.00446761 | 0.01263766 |
| 16 | 30 | 0 | 0.00541614 |
| 17 | 28 | 0.0014892 | 0.00469399 |
| 18 | 3 | 0 | 0.00054161 |
| 19 | 18 | 0 | 0.00324968 |
| 20 | 1 | 0 | 0.00018054 |
| 21 | 5 | 0 | 0.00090269 |
| 22 | 1 | 0 | 0.00018054 |

**Table S1. Detailed quantitative description of classes from low and high tau aggregation cells**

| **Class** | **Number of clusters per class** | **Ratio of Thr231 clusters over the total no of Thr231 clusters in each class** | **Ratio of AT8 clusters over the total no of AT8 clusters in each class** | **Ratio of GFP nano clusters over the total no of GFP nano clusters in each class** |
| --- | --- | --- | --- | --- |
| 1 | 1491 | 0.1822222 | 0.1217982 | 0.1768381 |
| 2 | 635 | 0.0355556 | 0.0893884 | 0.0662598 |
| 3 | 2318 | 0.4266667 | 0.2122321 | 0.2638768 |
| 4 | 358 | 0.0266667 | 0.0627287 | 0.0337111 |
| 5 | 98 | 0.0088889 | 0.0245687 | 0.00712 |
| 6 | 1025 | 0.1288889 | 0.1003659 | 0.1168265 |
| 7 | 609 | 0.0622222 | 0.0935703 | 0.0604475 |
| 8 | 132 | 0.0133333 | 0.026137 | 0.0114792 |
| 9 | 738 | 0.0933333 | 0.0773654 | 0.0826795 |
| 10 | 592 | 0.0177778 | 0.0794564 | 0.0633537 |
| 11 | 11 | 0 | 0.0010455 | 0.0013078 |
| 12 | 110 | 0.0044444 | 0.0130685 | 0.0122058 |
| 13 | 668 | 0 | 0.0737062 | 0.0765766 |
| 14 | 44 | 0 | 0.0078411 | 0.0042139 |
| 15 | 95 | 0 | 0.0115003 | 0.0106074 |
| 16 | 12 | 0 | 0.0005227 | 0.0015984 |
| 17 | 37 | 0 | 0.0036592 | 0.0043592 |
| 18 | 11 | 0 | 0.0005227 | 0.0014531 |
| 19 | 11 | 0 | 0 | 0.0015984 |
| 20 | 18 | 0 | 0.0005227 | 0.0024702 |
| 21 | 1 | 0 | 0 | 0.0001453 |
| 22 | 5 | 0 | 0 | 0.0007265 |
| 23 | 1 | 0 | 0 | 0.0001453 |

**Table S2.** **Detailed quantitative description of classes from Thr231, AT8 and GFP nanobody stained Clone 4.1 cells**

| **Antibody** | **Host species** | **Catalog number** | **Vendor** |
| --- | --- | --- | --- |
| α-tubulin | Rabbit | ab18251 | Abcam |
| T22 | Rabbit | ABN454 | Sigma-Aldrich |
| Thr231 | Mouse | MN1040 | ThermoFisher Scientific |
| AT8 | Mouse | MN1020 | ThermoFisher Scientific |
| GFP Binding Protein | Alpaca | gt-250 | ChromoTek |
| Tau-5 | Mouse | AHB0042 | ThermoFisher Scientific |

**Table S3. Antibodies used in this study**
